# Macrophage WDFY3, a protector against autoimmunity

**DOI:** 10.1101/2024.08.17.608411

**Authors:** Xun Wu, Ziyi Wang, Katherine R. Croce, Fang Li, Jian Cui, Vivette D. D’Agati, Rajesh K. Soni, Shareef Khalid, Danish Saleheen, Ira Tabas, Ai Yamamoto, Hanrui Zhang

**Affiliations:** Cardiometabolic Genomics Program, Division of Cardiology, Department of Medicine, Columbia University Irving Medical Center, New York, NY, USA; Department of Neurology, Columbia University, New York, NY, USA; Department of Pathology and Cell Biology, Columbia University, New York, NY, USA; Renal Pathology Laboratory, Columbia University Irving Medical Center, New York, NY, USA; Department of Medicine, Columbia University Irving Medical Center, New York, NY, USA; Department of Physiology and Cellular Biophysics, Columbia University Irving Medical Center, New York, NY, USA

**Keywords:** Macrophage, efferocytosis, inflammation, immunity

## Abstract

Efficient efferocytosis is essential for maintaining homeostasis. Excessive apoptotic cell (AC) death and impaired macrophage efferocytosis lead to autoantigen release and autoantibody production, immune activation, and organ damage. It remains unclear whether these immunogenic autoantigens are the sole cause of increased autoimmunity or if efferocytosis of ACs directly influences macrophage function, impacting their ability to activate T cells and potentially amplifying autoimmune responses. Additionally, it has not been established if enhancing macrophage efferocytosis or modulating macrophage responses to AC engulfment can be protective in autoimmune-like disorders. Our previous work showed WDFY3 is crucial for efficient macrophage efferocytosis. This study reveals that myeloid knockout of *Wdfy3* exacerbates autoimmunity in young mice with increased AC burden by systemic injections of ACs and in middle-aged mice developing spontaneous autoimmunity, whereas ectopic overexpression of WDFY3 suppresses autoimmunity in these models. Macrophages, as efferocytes, can activate T cells and the inflammasome upon engulfing ACs, which are suppressed by overexpressing WDFY3. This work uncovered the role of WDFY3 as a protector against autoimmunity by promoting macrophage efferocytosis thus limiting autoantigen production, as well as mitigating T cell activation and inflammasome activation.

## Main

Efficient efferocytosis of dying cells by phagocytes, mainly macrophages, is essential for maintaining homeostasis^1,2^. When the burden of apoptotic cells (ACs) exceeds the clearance capacity of efferocytosis, ACs can undergo necrosis and rupture, releasing cellular materials that become a continuous source of immunogenic stimuli, a well-known mechanism of autoimmune disorders^1,3^.

Excessive AC death and impaired efferocytosis are observed in patients with systemic lupus erythematosus^4–7^. Experimental studies establish that mice lacking genes involved in regulating efferocytosis have impaired clearance of ACs and develop lupus-like autoimmunity^8–12^, including genes coding for efferocytic receptors such as TIM4 or MERTK^8,9^, scavenging receptor SCARF1^10^, and soluble bridging proteins that recognize phosphatidylserine on ACs such as milk fat globule epidermal growth factor 8 (MFG-E8)^11,12^. However, while these whole-body knockout studies establish a role for defective efferocytosis in autoimmune-like phenotypes, it remains unclear whether myeloid-specific defects alone can drive systemic autoimmune responses. Open questions also remain about whether the immunogenic autoantigens produced due to insufficient efferocytosis are the sole cause of increased autoimmunity, or if efferocytosis of ACs directly impacts macrophage’s ability to activate T cells and potentially amplify autoimmune responses. Additionally, it has not been established if enhancing macrophage efferocytosis or modulating macrophage responses to AC engulfment can be protective in autoimmune-like disorders.

We have previously identified, via a genome-wide CRISPR screening, that WDFY3 is required for both the engulfment and the subsequent lysosomal acidification and degradation of ACs by macrophages^13^. Herein, we specifically eliminated *Wdfy3* expression in myeloid cells to determine whether defective macrophage efferocytosis drives autoimmunity in young mice with increased AC burden and in middle-aged mice that develop spontaneous autoimmunity. We then overexpress *WDFY3* to explore its therapeutic potential in suppressing autoimmune-related pathogenesis. We uncovered that, in addition to its role in regulating the efferocytic clearance of ACs, myeloid-specific loss of *Wdfy3* function exacerbates MHC-II mediated antigen presentation and inflammasome activation induced by AC engulfment. Moreover, ectopic overexpression of WDFY3 enhances efferocytosis, suppresses AC-induced inflammasome activation and T cell activation, and thereby mitigates autoimmunity. Our findings suggest that WDFY3 plays a pivotal role in mitigating autoimmune responses by enhancing efferocytosis thereby limiting self-antigen production, as well as mitigating efferocyte-T cell crosstalk and inflammasome activation.

### Myeloid knockout of *Wdfy3* exacerbates autoimmunity induced by systemic AC injections in young mice and spontaneous autoimmunity in middle-aged mice

Given our previous findings demonstrating the importance of WDFY3 in regulating multiple steps of the efferocytic clearance of ACs^13^, we set out to determine the pathological consequence of impaired efferocytosis in mice with myeloid knockout of *Wdfy3*. Delayed clearance of exogenously injected ACs leads to progressive development of autoantibodies^8^. Using a similar approach, we aimed to determine whether myeloid-specific knockout of *Wdfy3* will exacerbate autoimmune-like phenotypes induced by increasing the burden of ACs through systemic injections.

We backcrossed *Wdfy3^fl/fl^* mice^14^ generated by insertion of two *loxP* sites flanking exon 5 to mice on a C57BL/6J background for nine generations. Myeloid-specific *Wdfy3* null mice were obtained by breeding *Wdfy3^fl/fl^*mice with *LysMCre* mice for Cre-mediated recombination in the myeloid cell lineage^15^, including monocytes, mature macrophages, and granulocytes, i.e. *LysMCre^+/-^Wdfy3^fl/fl^*mice (Cre^+^), while using *LysMCre^-/-^Wdfy3^fl/fl^* littermates (Cre^-^) as the controls (as illustrated in **Fig. 1a**); and the loss of WDFY3 protein and impaired efferocytosis in Cre^+^ macrophages was confirmed^13^. Beginning at 8-10 weeks of age, Cre^-^ and Cre^+^ mice were injected with UV irradiation-induced apoptotic thymocytes over an 8-week period (**Fig. 1a**) and examined for markers of organ damage (**Fig. 1b,c**), serum autoantibodies (**Fig. 1d,e**), and kidney histology and immune complex deposition (**Fig. 1f-i**). Markers of organ damage, including serum alanine aminotransferase (ALT) and creatinine (**Fig. 1b,c**), were not different between Cre^-^ and Cre^+^ mice with AC injections. Cre^+^ mice that received AC injections showed higher serum antinuclear antibodies (ANA) compared to Cre^-^ mice with AC injections (**Fig. 1d**), indicating worsened autoimmune responses in Cre^+^ mice. Anti-dsDNAs (**Fig. 1e**), however, did not differ across genotypes. Cre^+^ mice exhibited more severe histological features of glomerulonephritis in response to AC injections, including enlarged area (**Fig. 1f**). H&E stain showed more glomerular mesangial hypercellularity in Cre^+^ mice compared to Cre^-^ mice with AC injections (**Fig. 1g**). Furthermore, Cre^+^ mice displayed increased C1q and IgG deposition in the glomerular mesangium compared to Cre^-^ mice with AC injections (**Fig. 1h,i**). The kidney immune complex deposition and nephritis-like phenotype were unlikely due to the direct effects of intrarenal AC accumulation, since the injected ACs mostly accumulated in the spleen (**Fig. S1**), suggesting that spleen is directly involved in the clearance of intravenously administered ACs. Rather, the increased AC burden and inefficient efferocytosis in Cre^+^ mice led to increased autoantigen-triggered autoantibody production and resulting immune complex-mediated glomerulonephritis.

**Fig. 1.**
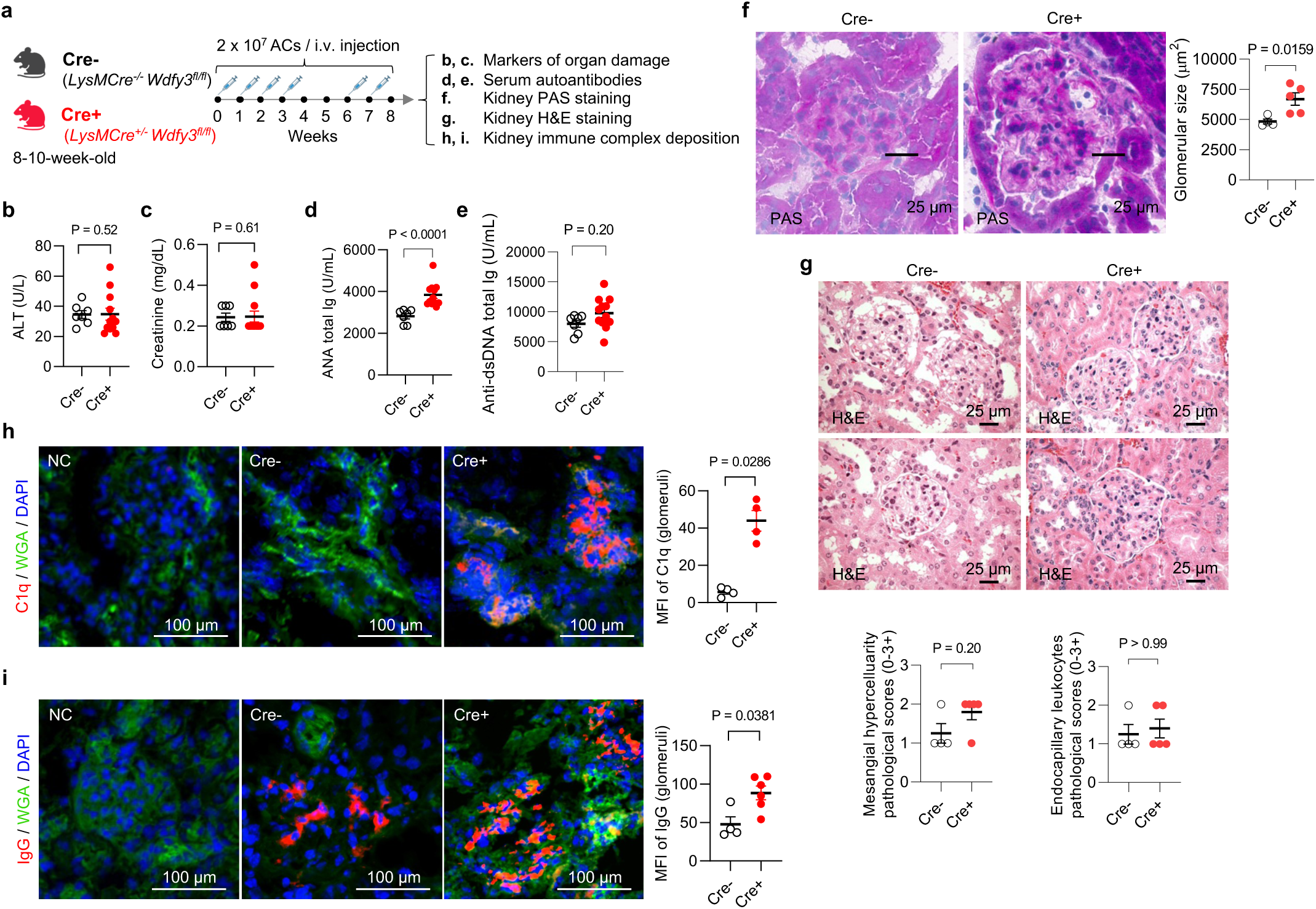
Myeloid knockout of *Wdfy3* exacerbates autoimmune responses in mice that receive apoptotic cell (AC) injections. **a,** Schematics of study design. Mice with *LysMCre-*mediated myeloid-specific knockout of *Wdfy3* (Cre^+^, *LysMCre^+/-^WDFY3^fl/fl^*) and the respective controls (Cre^-^, *LysMCre^-/-^WDFY3^fl/fl^*) were bred, and age- and sex-matched littermates were used for experiments. To increase the burden of apoptotic cells (ACs), murine thymocytes underwent UV irradiation to induce apoptosis. Subsequently, ACs were injected intravenously via the tail vein according to the timeline illustrated. Serum samples were collected to determine markers of organ damage and autoantibodies. Kidneys were dissected and sectioned for periodic acid-Schiff (PAS), H&E, and immunofluorescence staining. **b,** Serum alanine transaminase (ALT), a marker of liver damage. n = 7 Cre^-^ vs 12 Cre^+^ mice. **c**, Serum creatinine, a marker of kidney damage. n = 7 Cre^-^ vs 13 Cre^+^ mice. **d**, Serum anti-nuclear antibodies (ANA). n = 7 Cre^-^ vs 12 Cre^+^ mice. **e**, Serum anti-dsDNA antibodies. n = 7 Cre^-^ vs 12 Cre^+^ mice. **f**, PAS staining of kidney sections highlight glomerular basement membrane for the quantification of glomerular size. n = 4 Cre^-^ vs 5 Cre^+^ mice. **g**, H&E staining of kidney sections were performed to assess mesangial hypercellularity (left) and endocapillary hypercellularity/leukocyte infiltration (right). n = 4 Cre^-^ vs 5 Cre^+^ mice. **h** and **i,** Immunofluorescence staining of kidney sections using anti-C1q and anti-IgG antibodies was performed. Mean fluorescent intensity (MFI) in glomeruli were quantified. n = 4 Cre^-^ vs 4 Cre^+^ mice (**h**); n = 4 Cre^-^ vs 6 Cre^+^ mice (**i**). WGA, wheat germ agglutinin. Data are shown as mean ± SEM.

In young mice without AC burden, ALT, ANA, glomerular size or immune complex deposition in the kidney remain low and do not differ between Cre^-^ and Cre^+^ mice (**Fig. S2**), suggesting that myeloid knockout of *Wdfy3* was not sufficient to drive autoimmune-like disorders in young mice without an increased AC burden. At midlife (52-60-week-old), however, Cre^+^ mice demonstrated these autoimmune phenotypes in the absence of AC injections (**Fig. S3**), suggesting progressive autoimmune responses, likely due to increased cell death and change in immune cell function associated with age^16^. Consistent with what we have published in young mice, BMDMs of middle-aged Cre^+^ mice have a complete loss of WDFY3 protein (**Fig. S3b**) and impaired efferocytosis, including engulfment and acidification of the engulfed ACs, compared to Cre^-^ controls (**Fig. S3c**). Middle-aged Cre^+^ mice also showed increased spleen and kidney weight (**Fig. S3d**) and increased serum ALT, a marker of liver dysfunction (**Fig. S3e**), suggesting that chronic myeloid knockout of *Wdfy3* led to progressive organ damage. Creatinine was comparable between Cre^-^ and Cre^+^ mice (**Fig. S3f**), suggesting no functional kidney damage in these mice despite histological changes at this stage. Serum ANA was increased in Cre^+^ middle-aged mice (**Fig. S3g**), together with increased glomerular size (**Fig. S3i**) and increased C1q and IgG immune complex deposition in the glomeruli (**Fig. S3j**,**k**). Anti-dsDNA antibodies (**Fig. S3h**) were not different between middle-aged Cre^-^ and Cre^+^ mice. These observations suggest that myeloid knockout of *Wdfy3* causes a spontaneous, autoinflammatory, lupus-like syndrome in middle-aged mice, resembling the phenotypes observed in young mice with AC injections.

To explore human relevance, we queried 100,814 sequenced individuals from the Pakistan Genomic Resource (PGR), a cohort with a high rate of consanguinity in which rare loss-of-function (LOF) variants are more likely to be found^17,18^. We identified 13 carriers of a total of 9 putative LOF variants of *WDFY3*, including 5 frameshift and 4 essential splice variants. The low number of putative LOF variants is consistent with previous observations that putative LOF variants are depleted in this gene, and it is likely haplo-insufficient in humans^19^. The low number of putative LOF variant carriers precluded us from performing a well-powered LOF gene burden test. We therefore tested if missense variants affecting known functional domains of WDFY3^13,20^ are associated with autoimmune traits assessed in the PGR. We tested associations of missense variants with serum IL-18, IgG, ALT, AST (aspartate aminotransferase), and creatinine. Although the associations did not achieve significance at the 5% false discovery rate threshold, we observed nominal associations (p < 0.05). Specifically, carriers of missense variants exhibited increased levels of ALT, AST, and creatinine (**Table S1**). These findings are consistent with the increased ALT levels and kidney histology and immune complex deposition observed in middle-aged Cre^+^ mice compared to Cre^-^ mice (**Fig. S3**).

### Myeloid knockout of *Wdfy3* exacerbates autoimmunity via efferocytosis-dependent and macrophage-intrinsic mechanisms, including enhanced antigen presentation and inflammasome activation

Autoantigens released from dead cells that are not efficiently cleared by efferocytosis are processed and presented by antigen presenting cells, including macrophages, to CD4^+^ T cells that activate B cells to produce autoantibodies^21,22^. These autoantibodies deposit in organs, including kidney, leading to inflammation and pathological damages^23^. However, it remains unclear whether the autoimmune response is solely due to the increased release of autoantigen from dead cells because of impaired clearance, or if the cell-intrinsic effects within macrophages triggered by AC engulfment directly impact macrophages’ ability to activate T cells and potentially amplify autoimmune responses.

While myeloid knockout of *Wdfy3* was not sufficient to drive autoimmune-like disorders in young mice without an increased AC burden, we found that young mice with myeloid knockout of *Wdfy3* showed increased T cell activation (**Fig. 2b-g**). Specifically, Cre^+^ mice had an increased percentage of CD44^hi^CD62L^lo^ activated T cells in CD4^+^ T cells, though not in CD8^+^ T cells, in the blood (**Fig. 2b,c**). CXCR3, a chemokine receptor that is highly expressed on effector T cells^24^, was increased in both CD4^+^ and CD8^+^ T cells in the blood of Cre^+^ mice (**Fig. 2d,e**). The percentage of CD44^hi^CD62L^lo^ activated T cells in CD4^+^ T cells and CD8^+^ T cells were also increased in the spleen of Cre^+^ mice (**Fig. 2f,g**).

**Fig. 2.**
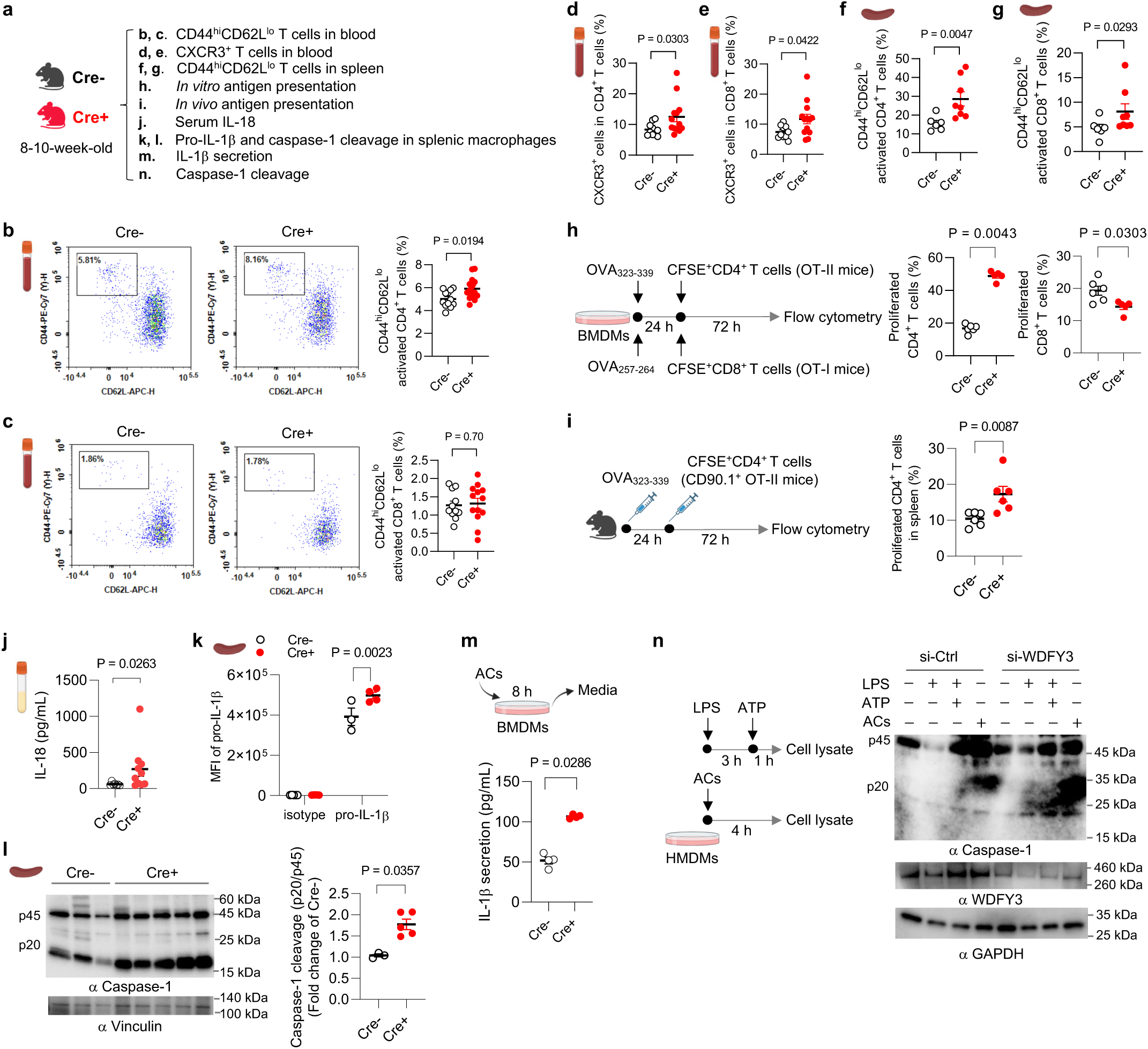
Myeloid knockout of *Wdfy3* augments T cell activation by promoting macrophage antigen presentation and inflammasome activation. **a,** Schematics of study design. Cre^-^ and Cre^+^ littermates were used to assess T cell activation (**b-g**), antigen presentation (**h** and **i**), and inflammasome activation (**j-n**). **b** and **c,** The percentage of CD44^high^CD62L^low^ activated T cells in CD4^+^ and CD8^+^ T cells in the blood. n = 12 Cre^-^ vs 14 Cre^+^ mice (**b**); n = 11 Cre^-^ vs 13 Cre^+^ mice (**c**). **d** and **e,** The percentage of CXCR3^+^ activated T cells in CD4^+^ and CD8^+^ T cells in the blood. n = 10 Cre^-^ vs 13 Cre^+^ mice. **f** and **g,** The percentage of CD44^high^CD62L^low^ activated T cells in splenic CD4^+^ and CD8^+^ T cells. n = 6 Cre^-^ vs 8 Cre^+^ mice. h, *In vitro* antigen presentation assay. BMDMs were pre-treated with synthetic ovalbumin (OVA) for 24 h, followed by incubating with splenic T cells isolated from OT-II or OT-I transgenic mice for 72 h to assess OVA_323-339_ or OVA_257-264_-specific activation of CD4^+^ T cells (OT-II mice) or CD8^+^ T cells (OT-I mice) that recognize OVA peptide residues when presented by the MHC-II or MHC-I molecules, respectively. The OT-II or OT-I mouse T cells were pre-labeled by carboxy-fluorescein succinimidyl ester (CFSE) to monitor the dilution of CFSE signal in daughter cells as T cells become activated and divide. n = 6 Cre^-^ vs 5 Cre^+^ mice. **i**, *In vivo* antigen presentation assay. Mice were immunized with OVA_323-339_ for 24 h before the injection of CFSE-labeled splenic CD4^+^ T cells from CD90.1^+^ OT-II mice. 72 h after injection, CFSE dilution in CD4^+^CD90.1^+^ splenic T cells was assessed. n = 6 Cre^-^ vs 6 Cre^+^ mice. **j,** Serum IL-18. n = 7 Cre^-^ vs 10 Cre^+^ mice. **k,** The expression of pro-IL-1β in F4/80^+^ splenic macrophages assessed by flow cytometry. n = 5 Cre^-^ vs 5 Cre^+^ mice in isotype group; n = 3 Cre^-^ vs 4 Cre^+^ mice in pro-IL-1β group. **l,** Analysis of intracellular caspase-1 cleavage in F4/80+ splenic macrophages by Western blotting. The intensity of the active form of caspase-1, p20, was quantified and compared to the intensity of the pro-caspase-1 (p45). The ratio of p20 intensity to p45 intensity was calculated, and this ratio was normalized to the control group (Cre^-^), which was assigned a value of 1.0 for standardization. n = 3 Cre^-^ vs 5 Cre^+^ mice. **m,** BMDMs were co-incubated with ACs at a 5:1 AC-to-BMDM ratio for 8 h. Culture media were collected to determine IL-1β by ELISA. n = 4 Cre^-^ vs 4 Cre^+^ mice. **n,** Human peripheral blood monocytes were differentiated into macrophages (HMDMs) followed by transfection of non-targeting control siRNA (si-Ctrl) or siRNAs targeting *WDFY3* (si-WDFY3). At day 7, HMDMs were primed with 20 ng/mL lipopolysaccharide (LPS) for 3 h and then treated with 2 mM ATP for an additional 1 h to induce inflammasome activation. To assess if AC induces caspase-1 cleavage, we co-incubated ACs with HMDMs for 4 h at a 5:1 AC-to-HMDM ratio. Unbound ACs were washed away. HMDMs were harvested to determine caspase-1 cleavage by Western blotting. The representative blot from three technical replicates was shown. Data are shown as mean ± SEM.

T cells are activated when their receptors recognize antigens presented by antigen-presenting cells, such as macrophages, dendritic cells, and B cells, and receive additional signals from cytokines^25–27^. To determine if macrophage WDFY3 directly regulates antigen presentation capacity by macrophages via cell intrinsic mechanisms, we performed a standard antigen presentation assay using soluble antigens, which require the interaction of the MHC complex on ovalbumin (OVA)-treated macrophages with T cells isolated from OT-II or OT-I transgenic mice (**Fig. 2h**). The OT-II transgenic mice express mouse T cell receptors designed to specifically recognize OVA peptide residues 323-339 (OVA_323-339_), when presented to CD4^+^ T cells by the MHC-II molecule, whereas OT-I transgenic mice express mouse T cell receptors designed to recognize OVA peptide residues 257-264 (OVA_257-264_), when presented to the CD8^+^ T cells by the MHC-I molecule. We find that knockout of *Wdfy3* in macrophages leads to increased MHC-II mediated antigen presentation to CD4^+^ T cells, while decreased MHC-I mediated antigen presentation to CD8^+^ T cells (**Fig. 2h**)

To confirm our findings *in vivo*, we immunized Cre^+^ mice and their control counterparts with OVA_323-339_, followed by an adoptive transplantation of CD4^+^ T cells from CD90.1^+^ OT-II mice^25^, which specifically recognize OVA-MHC-II complexes. The distinction of donor T cells from the recipient’s T cells relies on the expression of CD90.1, a surface marker that is unique to the donor T cells, while the recipient mice express CD90.2^25^. To monitor proliferation, these CD4^+^ CD90.1^+^ OT-II T cells were labeled with carboxyfluorescein succinimidyl ester (CFSE). In Cre^+^ mice, there was a significant CFSE dilution in CD4^+^ CD90.1^+^OT-II T cells in the spleen (**Fig. 2i**), indicating enhanced antigen presentation by *Wdfy3*-deficient splenic macrophages that promotes T cell proliferation. The results suggest that loss of WDFY3 regulates MHC-II mediated antigen presentation to the surface of macrophage, contributing to increased antigen presentation to CD4^+^ T cells and enhanced activation. On the other hand, the activation of CD8^+^ T cells observed in young Cre^+^ mice *in vivo* was unlikely to be directly mediated by macrophage-intrinsic mechanisms, as knockout of *Wdfy3* in macrophages resulted in an decrease in MHC-I mediated antigen presentation to CD8^+^ T cells *in vitro* (**Fig. 2h**).

We next determined if T cell activation is associated with increased systemic inflammatory cytokine levels. We used a 26-Plex ProcartaPlex mouse cytokine and chemokine panel, which includes cytokines produced by activated T cells or those that instruct T cell activation (**Fig. S4**). Through this initial screening, we found that only IL-18 was increased in the serum of Cre^+^ mice (**Fig. 2j**). IL-18, together with IL-1β and IL-1α, are pro-inflammatory cytokines in the IL-1 family^28,29^. Their secretion, especially the secretion of IL-1β and IL-18, is regulated by inflammasomes which are multi-protein complexes containing sensor proteins, adaptor protein and caspase-1^28,29^. IL-1β and IL-18 are the direct indicator of inflammasome activation and caspase-1 is a key effector enzyme involved in the cleavage and activation of pro-IL-1β and pro-IL-18. We then determined whether the knockout *Wdfy3* in macrophages affects inflammasome activation. Indeed, splenic F4/80^+^ macrophages of Cre^+^ mice show increased pro-IL-1β protein expression (**Fig. 2k**) and caspase-1 cleavage (**Fig. 2l**), suggesting enhanced priming and activation of the inflammasome.

To test whether *Wdfy3* knockout in macrophages regulates inflammasome activation in the context of efferocytosis, we determined IL-1β secretion in BMDMs co-incubated with ACs for 8 h. *Wdfy3*-deficient BMDMs showed increased IL-1β secretion (**Fig. 2m**). To establish human relevance, we determined the effects of siRNA-mediated *WDFY3* knockdown in human monocyte-derived macrophages (HMDMs). Either treatment with ACs, or LPS and ATP that activate the NLRP3 inflammasome, led to increased caspase-1 cleavage in HMDMs (**Fig. 2n**). *WDFY3* knockdown further exacerbates caspase-1 cleavage (**Fig. 2n**).

IL-1β is known to enhance antigen-primed CD4^+^ and CD8^+^ T-cell activation^30^. IL-18 and 1L-12 synergize for IFN-γ production in Th1 cells^31^. We, however, did not observe increased IL-12 or IFN-γ (**Fig. S4**), suggesting that IL-18 did not directly contribute to T cell activation in young myeloid-specific *Wdfy3* knockout mice. Taken together, myeloid *Wdfy3* knockout leads to increased T cell activation likely via efferocytosis-dependent and macrophage-intrinsic mechanisms by enhancing antigen presentation capacity and inflammasome activation.

### Overexpression of WDFY3 enhances macrophage efferocytosis *in vitro* and *in vivo*

Given how the loss of WDFY3 diminishes efferocytosis and promotes autoimmunity, we next investigated whether increasing expression levels of WDFY3 could enhance efferocytosis and reduce autoimmunity. Mice for overexpressing human *WDFY3* were generated by knocking the construct, *loxP-STOP-loxP-3xFLAG-hWDFY3 cDNA*, into the *Rosa26* locus (*Rosa26^loxP-STOP-loxP-3xFLAG-hWDFY3^ ^cDNA/+^*)^32^. To achieve constitutive overexpression, male mice carrying the floxed alleles were crossed with female carrier of *HprtCre*^33^, a Cre-deleter strain with 100% Cre-mediated recombination in oocytes to generate heritable deletion of *loxP-STOP-loxP* cassette (*Rosa26^3xFLAG-hWDFY3/+^*). *Rosa26^3xFLAG-hWDFY3/+^* mice were interbred to generate homozygotes knock-in (KI) mice (hWDFY3^KI/KI^) and wildtype littermate controls (Ctrl) and were maintained on a mixed background (C57BL/6 x 129/SvEv).

BMDMs from hWDFY3^KI/KI^ mice show higher levels of WDFY3 expression by Western blotting (**Fig. 3b**), accompanied by higher engulfment capacity compared to their control counterparts (**Fig. 3c**). Consistent with our loss of function studies showing impaired LC3 lipidation and lysosomal acidification of engulfed ACs in macrophages of Cre^+^ mice^13^, BMDMs of WDFY3-overexpressing mice had a modest increase in the percentage of engulfed ACs that were also acidified (**Fig. 3c**). This was accompanied by increased LC3 lipidation (**Fig. 3d**), the final step required for phagosome-lysosome fusion and the subsequent lysosomal acidification^34^. This modest increase suggests that the enhanced engulfment of cargo is associated with efficient degradation. We also confirmed enhanced efferocytosis in peritoneal macrophages (PMs) of WDFY3-overexpressing mice (**Fig. 3e**).

**Fig. 3.**
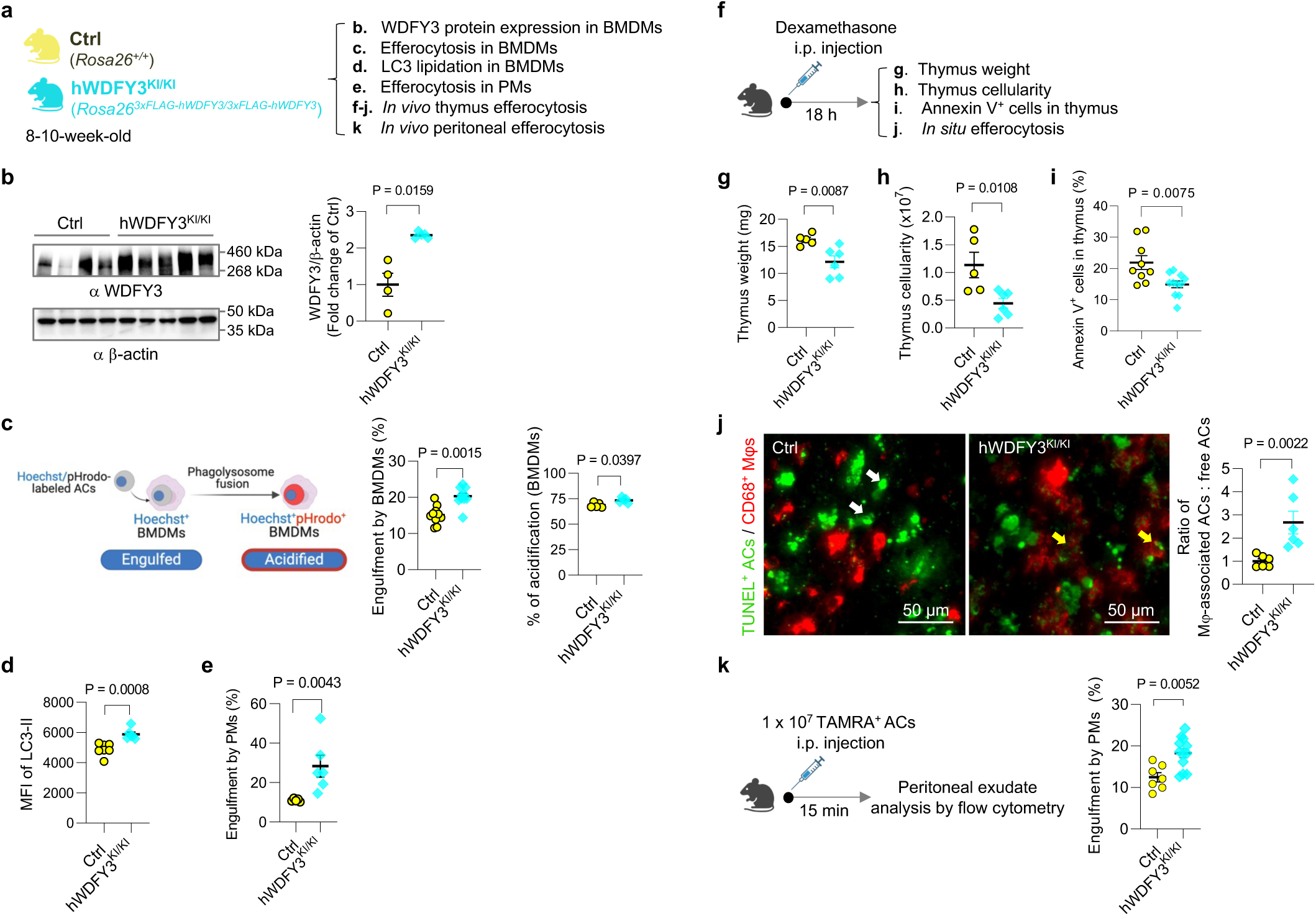
Overexpression of *WDFY3* enhances efferocytosis *in vitro* in macrophages and *in vivo* in mice. **a,** Schematics of experimental design. Mice with ectopic overexpression of human *WDFY3* were generated by knock-in (KI) of the construct, i.e., cDNA sequence encoding 3x FLAG tag that is fused to human *WDFY3* cDNA preceded by a STOP cassette flanked by *loxP* sites, into the *Rosa26* locus (*Rosa26^loxP-STOP-loxP-3xFLAG-hWDFY3/+^*). Constitutive overexpression was achieved by breeding male mice carrying the floxed allele with female *HprtCre* to generate heritable deletion of the *loxP-STOP-loxP* cassette (*Rosa26^3xFLAG-hWDFY3/+^*). *Rosa26^3xFLAG-hWDFY3/+^*mice were interbred to generate homozygotes *Rosa26^3xFLAG-hWDFY3/3xFLAG-hWDFY3^*(hWDFY3^KI/KI^) and wildtype littermate controls *Rosa26^+/+^* (Ctrl). **b**, Validation of overexpression of WDFY3 in BMDMs by Western blotting. n = 4 Ctrl vs 5 hWDFY3^KI/KI^ mice. **c,** BMDMs were incubated with ACs labeled by Hoechst, which stains DNA and is pH-insensitive, and pHrodo-Red, which is pH-sensitive and fluoresces only in an acidified environment in the phagolysosome. The percentage of Hoechst^+^ BMDMs indicates engulfment. n = 10 Ctrl vs 10 hWDFY3^KI/KI^ mice. The percentage of Hoechst^+^/pHrodo^+^ BMDMs in Hoechst^+^ BMDMs indicates acidification of the engulfed corpses. n = 5 Ctrl vs 5 hWDFY3^KI/KI^ mice. **d,** BMDMs were incubated with Hoechst-labeled ACs to allow for efferocytosis. After removal of unbound ACs, BMDMs were collected and treated with digitonin to remove non-membrane bound LC3, and then immunostained for LC3 that is lipidated and membrane-bound, i.e. LC3-II. LC3-II staining was then quantified by flow cytometry for BMDMs that had engulfed Hoechst-labeled ACs. n = 7 Ctrl vs 6 hWDFY3^KI/KI^ mice. **e,** Peritoneal exudates were seeded and nonadherent cells were removed. The remaining peritoneal macrophages (PMs) were cultured for 18 h and then incubated with Hoechst-labeled ACs to allow for efferocytosis. After 1 h, unbound ACs were washed away, and PMs were harvested to quantify the % of Hoechst^+^ PMs by flow cytometry. n = 5 Ctrl vs 6 hWDFY3^KI/KI^ mice. **f,** Schematics of experimental design for *in vivo* thymus efferocytosis. Mice were injected intraperitoneally with dexamethasone to induce thymocyte apoptosis. 18 h after injection, thymi were dissected for assessment of the following parameters. **g**, Thymus weight. n = 5 Ctrl vs 6 hWDFY3^KI/KI^ mice. **h,** Total number of cells per thymus. n = 5 Ctrl vs 6 hWDFY3^KI/KI^ mice. **i,** The percentage of Annexin V^+^ cells per thymus. n = 9 Ctrl vs 11 hWDFY3^KI/KI^ mice. **j**. *In situ* efferocytosis. Thymic sections were stained with TUNEL for ACs, and CD68 for macrophages. The ratio of macrophage-associated TUNEL^+^ cells vs. free TUNEL^+^ cells was quantified to assess *in situ* efferocytosis in the thymus. The white and yellow arrows highlight free and macrophage-associated TUNEL^+^ cells, respectively. Lower weight and cellularity, lower percentage of Annexin V^+^ cells, and higher *in situ* efferocytosis indicate enhanced efferocytosis capacity. n = 6 Cre^-^ vs 6 hWDFY3^KI/KI^ mice. **k,** *In vivo* PM efferocytosis assay. 1 × 10^7^ TAMRA^+^ ACs were injected intraperitoneally, and peritoneal exudates were collected after 15 min. After blocking with CD16/32, peritoneal cells were stained with F4/80 antibody to label PMs and the percentage of TAMRA^+^ PMs was determined by flow cytometry. n = 7 Ctrl vs 12 hWDFY3^KI/KI^ mice. Data are shown as mean ± SEM.

We further validated efferocytosis in mice using two *in vivo* assays: thymus efferocytosis and PM efferocytosis^13^. The *in vivo* thymus efferocytosis assay involved injecting dexamethasone intraperitoneally into Ctrl and hWDFY3^KI/KI^ mice to induce thymocyte apoptosis (**Fig. 3f**), as the thymus is particularly sensitive to dexamethasone-induced apoptosis^35^. After 18 hours, we assessed thymus weight and cellularity (**Fig. 3g,h**). Compared to the Ctrl mice, hWDFY3^KI/KI^ mice showed significantly lower thymus weights and decreased cellularity (**Fig. 3g,h**), indicating enhanced efferocytic clearance of apoptotic thymocytes. Consistently, thymi from hWDFY3^KI/KI^ mice also contained fewer Annexin V^+^ cells (**Fig. 3i**). We next assessed *in situ* efferocytosis using TUNEL staining on thymus sections to calculate the ratio of TUNEL^+^ cells associate with CD68^+^ macrophages and free TUNEL^+^ cells not in proximity to macrophages (**Fig. 3j**). The data showed an increased ratio of macrophage-associated AC: free AC in hWDFY3^KI/KI^ mice, indicating increased efferocytosis in the thymus. We further confirmed our findings by assessing PM efferocytosis *in vivo*. Peritoneal exudate was collected 15 min after injecting TAMRA-labeled ACs^13^. PMs from hWDFY3^KI/KI^ mice showed an increased percentage of TAMRA-labeled F4/80^+^ macrophages, indicating enhanced efferocytotic capacity in the hWDFY3^KI/KI^ mice (**Fig. 3k**).

### Overexpression of WDFY3 attenuates autoimmunity induced by systemic AC injections in young mice and spontaneous autoimmunity in middle-aged mice

After confirming the enhanced efferocytic capacity in macrophages overexpressing WDFY3, we asked whether WDFY3 overexpression in mice provides protection against autoimmunity. We challenged hWDFY3^KI/KI^ mice with repeated injections of ACs as described for loss of function studies in **Fig. 1** and as illustrated in **Fig. 4a**, and assessed organ damage markers, circulating autoantibodies, kidney histology and immunocomplex deposition (**Fig. 4**). Organ damage markers remain low and comparable between Ctrl and hWDFY3^KI/KI^ mice (**Fig. 4b,c**). However, compared to Ctrl mice, hWDFY3^KI/KI^ mice showed lower levels of ANA and ant-dsDNA (**Fig. 4,e**), smaller glomeruli (**Fig. 4f**), and reduced C1q and IgG deposition in the kidney (**Fig. 4g,h**), indicating a protective effect from AC injections.

**Fig. 4.**
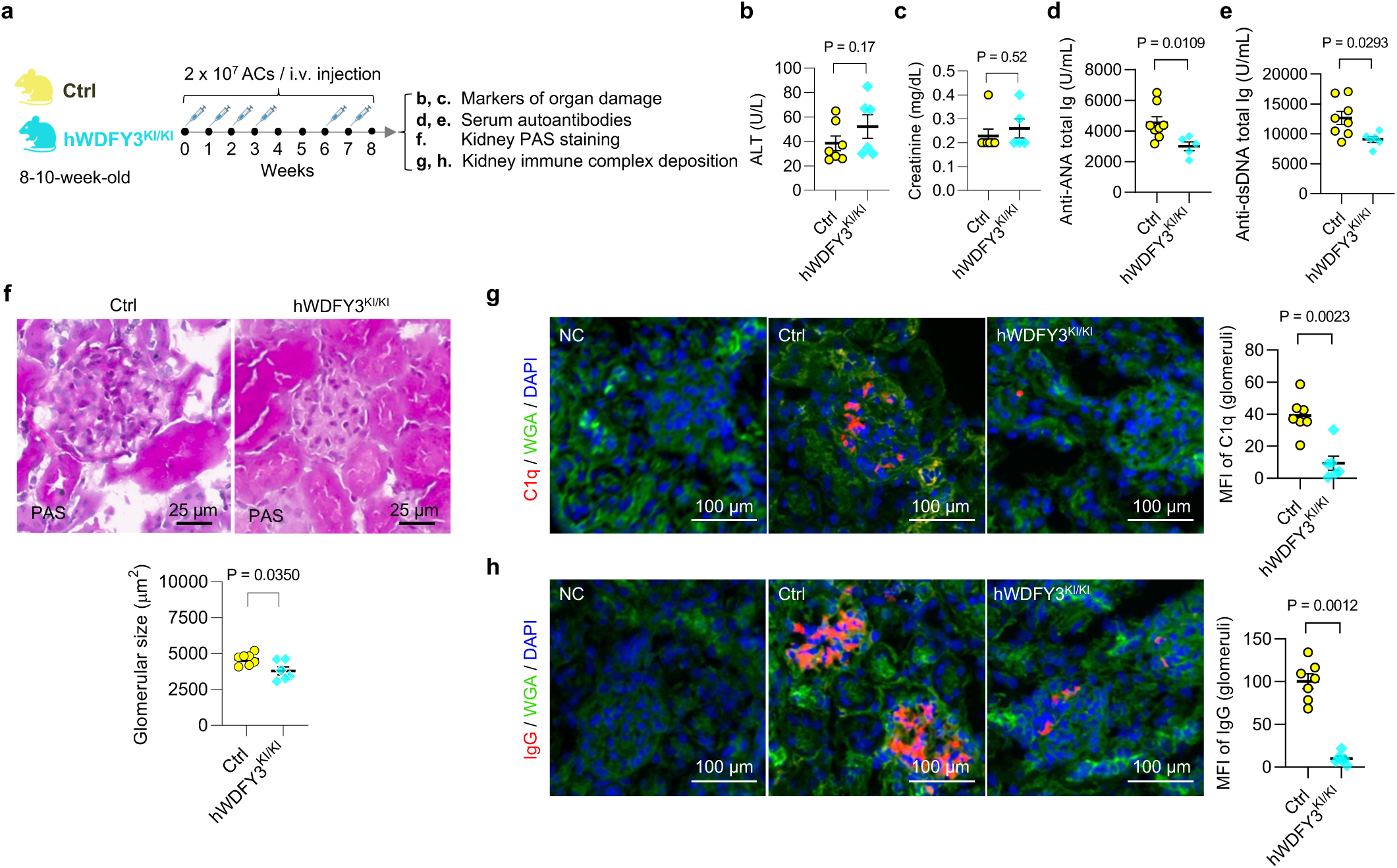
Overexpression of *WDFY3* mitigates autoimmune responses in mice that receive AC injections. **a.** Schematics of study design. Age- and sex-matched Ctrl and hWDFY3^KI/KI^ littermates were used for experiments. ACs were prepared and injected as illustrated and as described in Fig. 1a. Serum samples were collected to determine markers of organ damage and autoantibodies. Kidneys were dissected and sectioned for PAS and immunofluorescence staining. **b,** Serum ALT. n = 7 Ctrl vs 6 hWDFY3^KI/KI^ mice. **c**, Serum creatinine. n = 7 Ctrl vs 5 hWDFY3^KI/KI^ mice. **d**, Serum ANA. n = 8 Ctrl vs 5 hWDFY3^KI/KI^ mice. **e**, Serum anti-dsDNA antibodies. n = 8 Ctrl vs 6 hWDFY3^KI/KI^ mice. **f**, PAS staining of kidney sections for the quantification of glomerular size. n = 7 Ctrl vs 6 hWDFY3^KI/KI^ mice. **g** and **h,** Immunofluorescence staining of kidney sections using anti-C1q and anti-IgG antibodies. n = 7 Ctrl vs 6 hWDFY3^KI/KI^ mice. Data are shown as mean ± SEM.

The protective effects of WDFY3 overexpression also extends to middle-aged mice without AC injections (**Fig. S5**). Middle-aged hWDFY3^KI/KI^ mice showed enhanced macrophage efferocytosis (**Fig. S5b**), lower kidney and lymph node (LN) weight (**Fig. S5c**), lower ALT levels (**Fig. S5d**), and reduced ANA and anti-dsDNA (**Fig. S5f**,**g**), compared to middle-aged Ctrl mice, while creatinine level remained low and comparable (**Fig. S5e**). Additionally, hWDFY3^KI/KI^ mice exhibited smaller glomeruli (**Fig. S5h**) and lower IgG deposition in the kidney (**Fig. S5i**).

Collectively, these findings highlight the potential of WDFY3 overexpression as a therapeutic strategy for mitigating autoimmunity caused by increased dead cell burden and age-associated chronic inflammation.

### Overexpression of WDFY3 in macrophages mitigates T cell activation and inflammasome activation in response to the efferocytosis of ACs

We next examined whether WDFY3 overexpression suppresses T cell activation and inflammasome activation (**Fig. 5**). The percentage of CD44^hi^CD62L^lo^ activated T cells in CD4^+^ and CD8^+^ T cells, as well as CXCR3^+^ activated T cells in CD4^+^ and CD8^+^ T cells in the blood, showed a modest reduction in hWDFY3^KI/KI^ mice (**Fig. 5b-e**). The percentage of CD44^hi^CD62L^lo^ activated T cells in splenic CD4^+^ and CD8^+^ T cells also decreased in hWDFY3^KI/KI^ mice (**Fig. 5f,g**). We designed an experiment to directly assess efferocyte-T cell interaction based on a previous published protocol with modification^36^. ACs were co-cultured with BMDMs for 24 h to induce efferocytosis and then washed away, followed by the addition of CFSE-labeled CD4^+^ T cells to assess T cell proliferation over 72 h. Efferocytic BMDMs induced T cell activation, which was attenuated in BMDMs from hWDFY3^KI/KI^ mice (**Fig. 5h**). Moreover, hWDFY3^KI/KI^ mice showed enhanced MHC-II mediated antigen presentation to CD4^+^ T cells *in vivo* (**Fig. 5i**). Taken together, WDFY3 overexpression suppresses MHC-II mediated antigen presentation and efferocyte-T cell interaction, thereby attenuating T cell activation.

**Fig. 5.**
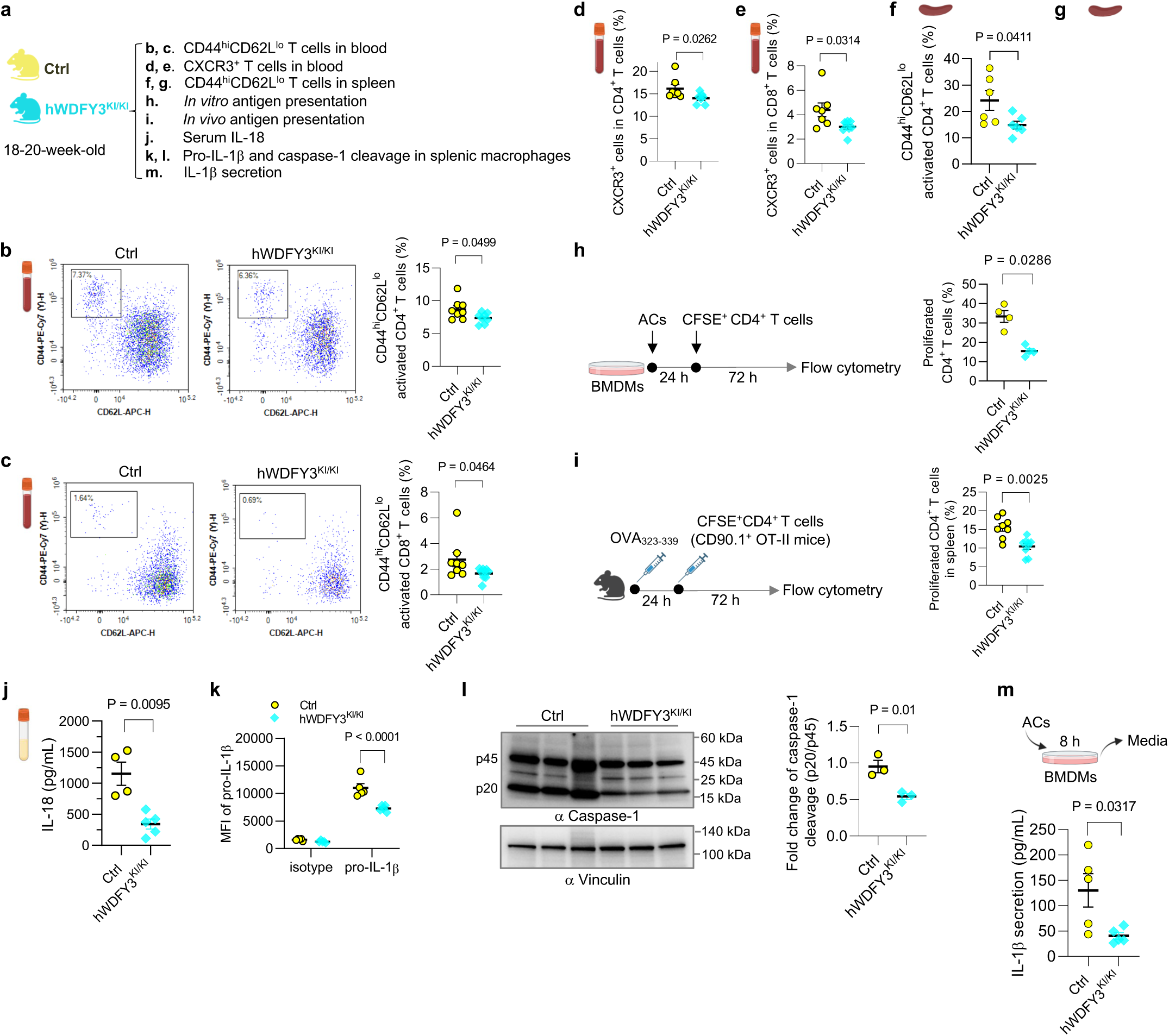
Overexpression of *WDFY3* suppresses T cell activation by restraining macrophage antigen presentation and inflammasome activation. **a,** Schematics of study design. Ctrl and hWDFY3^KI/KI^ littermates were used to assess T cell activation (**b-g**), antigen presentation (**h** and **i**), and inflammasome activation (**j-m**). **b** and **c,** The percentage of CD44^high^CD62L^low^ activated T cells in CD4^+^ and CD8^+^ T cells in the blood. n = 8 Ctrl vs 8 hWDFY3^KI/KI^ mice (**b**); n = 8 Ctrl vs 9 hWDFY3^KI/KI^ mice (**c**). **d** and **e,** The percentage of CXCR3^+^ activated T cells in CD4^+^ and CD8^+^ T cells in the blood. n = 7 Ctrl vs 7 hWDFY3^KI/KI^ mice (**d**); n = 7 Ctrl vs 8 hWDFY3^KI/KI^ mice (**e**). **f** and **g,** The percentage of CD44^high^CD62L^low^ activated T cells in splenic CD4^+^ and CD8^+^ T cells. n = 6 Ctrl vs 6 hWDFY3^KI/KI^ mice (**f**); n = 5 Ctrl vs 5 hWDFY3^KI/KI^ mice (**g**). **h,** Efferocyte-T cell interaction. BMDMs were pre-incubated with ACs for 24 h to induced efferocytosis, followed by incubating with CFSE-labeled CD4^+^ splenic T cells for 72 h to assess efferocyte-mediated activation of CD4^+^ T cells. n = 4 Ctrl vs 4 hWDFY3^KI/KI^ mice. **i**, *In vivo* antigen presentation assay. Mice were immunized with OVA323-339 for 24 h before the injection of CFSE-labeled splenic CD4+ T cells from CD90.1^+^ OT-II mice. 72 h after injection, CFSE dilution in CD4^+^CD90.1^+^ splenic T cells was assessed. n = 8 Ctrl vs 9 hWDFY3^KI/KI^ mice. **j,** Serum IL-18 in 15-month-old mice. n = 4 Ctrl vs 5 hWDFY3^KI/KI^ mice. **k,** The expression of pro-IL-1β in F4/80^+^ splenic macrophages assessed by flow cytometry. n = 5 Ctrl vs 5 hWDFY3^KI/KI^ mice in isotype group; n = 5 Ctrl vs 5 hWDFY3^KI/KI^ mice in pro-IL-1β group. **l,** Analysis of intracellular caspase-1 cleavage in F4/80^+^ splenic macrophages by Western blotting. The ratio of p20 intensity to p45 intensity was calculated, and this ratio was normalized to the control group (Ctrl), which was assigned a value of 1.0 for standardization. n = 3 Ctrl vs 3 hWDFY3^KI/KI^ mice. **m,** BMDMs were co-incubated with ACs at a 5:1 AC-to-BMDM ratio for 8 h. Culture media were collected to determine IL-1β by ELISA. n = 5 Ctrl vs 5 hWDFY3^KI^.

We also observed lower serum IL-18 levels in 15-month-old hWDFY3^KI/KI^ mice compared to Ctrl (**Fig. 5j**). The hWDFY3^KI/KI^ mice showed decreased pro-IL-1β expression and caspase-1 cleavage in splenic macrophages (**Fig. 5k,l**). Additionally, AC-induced IL-1β secretion was lower in hWDFY3^KI/KI^ BMDMs (**Fig. 5m**), suggesting that WDFY3 overexpression suppressed AC-induced inflammasome activation.

We speculate that the molecular mechanisms of WDFY3 function involve its role as an adaptor protein in autophagy-mediated protein aggregate degradation^37–40^. This hypothesis emerges in part from RNA-seq analysis of BMDMs from Ctrl and hWDFY3^KI/KI^ mice, which revealed only 13 differentially expressed genes (**Table S2**), suggesting minimal effects of WDFY3 overexpression at the transcriptomic level. Given the critical role of macroautophagy-dependent degradation in the regulation of NLRP3 inflammasome activation^41^, WDFY3 may suppress inflammasome activation by promoting its degradation. Additionally, WDFY3 localizes to RAB5- and EEA1-positive early endosomes^42^, where it may regulate antigen processing and peptide loading to MHC-II^43,44^. It is worth highlighting that successful efferocytosis, in particular the degradation of the AC corpses, induces the production of anti-inflammatory cytokines such as transforming growth factor-beta (TGF-β) and interleukin-10 (IL-10), promoting the resolution of inflammation^45–48^. Our work reveals that macrophage responses to AC engulfment involve inflammasome activation and T cell activation, which likely represent the initial responses. These macrophage responses are suppressed by WDFY3. Altogether, we uncovered new roles of macrophage WDFY3 in enhancing efferocytosis and suppressing antigen presentation and inflammasome activation, further augmenting the inflammation-resolving benefits of macrophage efferocytosis.

## Discussion

Our work advances several conceptual ideas on macrophage efferocytosis in autoimmunity. First, we identified a significant role for the myeloid-specific knockout of *Wdfy3* in promoting autoimmunity. This occurs in young mice with an increased AC burden and in middle-aged mice developing spontaneous autoimmunity. The underlying mechanisms involve impaired clearance of dead cells by efferocytosis, leading to autoantigen-triggered autoantibody formation and immune complex deposition, most notably in the kidney. Mechanistically, macrophages, as the efferocytes, can activate T cells and the inflammasome upon engulfing apoptotic cells, suggesting that the process of efferocytosis directly impact macrophage function to potentially amplify autoimmune responses. WDFY3 plays a critical role in regulating macrophage-T cell interaction by suppressing MHC-II mediated antigen presentation and inflammasome activation. Finally, overexpressing WDFY3 further enhances macrophage efferocytosis, mitigates AC-induced inflammasome activation, and suppresses T cell activation. Consequently, WDFY3 overexpression protects mice from autoimmune-like phenotypes triggered by AC injections as well as spontaneous, progressive autoimmunity developed in middle-aged mice.

Among immune cells, WDFY3 is abundantly expressed in monocytes and macrophages, with low expression in granulocytes, B cells and dendritic cells^13^. Beyond immune cells, WDFY3 is most abundantly expressed in neuronal cells, where it regulates the clearance of aggregated proteins through aggrephagy^38,40^. Ectopic overexpression of WDFY3 exerts neuroprotective effects in mice^32^. Although our work using constitutive whole-body overexpression mice does not exclude potential protective effects from other cell types, the essential role of macrophage WDFY3 as a protector against autoimmunity is well established using *in vitro* cultured murine and human macrophages and *in vivo* mouse models with myeloid-specific knockout. Our findings highlight the potential therapeutic applications of enhancing WDFY3 to mitigate autoimmune disorders associated with defective efferocytosis, as well as age-associated chronic inflammation^49^.

## Methods

### Mice

Animal protocols were approved by the Institutional Animal Care and Use Committee at Columbia University (Protocol Number AC-AABN5558). Mice were housed socially in standard cages at 22°C with 40-60% humidity under a 12-12 h light-dark cycle. *Ad libitum* access to water and food is provided by the mouse barrier facility (PicoLab Rodent Diet 20 5053 and 5058, LabDiet). Myeloid-specific *Wdfy3* knockout mice were generated by crossing *Wdfy3^fl/fl^* (generated Dr. Ai Yamamoto’s lab^14^ and backcrossed to C57BL/6J mice for nine generations) with *LysMCre^+/-^*(Stock No. 004781, the Jackson Laboratory)^15^. *LysMCre^+/-^Wdfy3^fl/fl^* mice (Cre^+^) had myeloid-specific knockout of *Wdfy3*, while *LysMCre^-/-^Wdfy3^fl/fl^* littermates (Cre^−^) served as controls^13^. Mice overexpressing human *WDFY3* were generated by Dr. Ai Yamamoto’s lab^32^. Specifically, the construct (the cDNA sequence encoding 3x FLAG tag that is fused to human *WDFY3* cDNA preceded by a STOP cassette flanked by *loxP* sites) was knocked into the *Rosa26* locus (*Rosa26^loxP-STOP-loxP-3xFLAG-hWDFY3^*^/^*^+^*) and the expression driven by the endogenous *Rosa26* promoter. To achieve constitutive overexpression, male mice carrying the floxed allele were crossed with female carriers of *HprtCre*, a Cre-deleter strain with 100% Cre-mediated recombination in oocytes (Stock No. 004302, the Jackson Laboratory) to generate heritable deletion of *loxP-STOP-loxP* cassette (*Rosa26^3xFLAG-hWDFY3/+^*). *Rosa26^3xFLAG-hWDFY3/+^* mice were interbred to generate homozygotes, *Rosa26^3xFLAG-hWDFY3/3xFLAG-hWDFY3^*, for knock-in (KI) of *3xFLAG-hWDFY3* (hWDFY3^KI/KI^) and wildtype littermate controls (Ctrl). The hWDFY3^KI/KI^ mice are on a mixed background (81.3% C57BL/6 and 18.7% 129/SvEv). The OT-I mice (Stock No. 003831)^50^, OT-II mice (Stock No. 004149)^51^, and CD90.1 (Thy 1.1) mice (Stock No. 000406) were obtained from the Jackson Laboratory.

### Macrophage isolation and differentiation

#### Murine bone marrow-derived macrophages (BMDMs)

Murine bone marrow-derived macrophages (BMDMs) were prepared by differentiating mouse bone marrow (BM) cells isolated from femurs and tibia^13^. BM cells were plated in 6-well non-tissue-culture (TC)-treated plates at 2 x 10^6^ to 3 x 10^6^ cells per well in DMEM basal media supplemented with 10% heat-inactivated (HI)- FBS, 20% L-929 fibroblast conditioned media, and 2 mM L-Glutamine^13^. BM cells are fully differentiated to BMDMs by day 7 for *in vitro* assay^13^.

#### Murine peritoneal macrophages (PMs)

Mouse peritoneal macrophages (PMs) were prepared by collecting peritoneal exudates. The exudates were centrifuged at 400 x *g* for 5 min to obtain cell pellets, which were resuspended and plated in 12-well non-TC-treated plates at 0.5 x 10^6^ cells per well in DMEM basal media supplemented with 10% heat-inactivated (HI)- FBS, 20% L-929 fibroblast conditioned media, and 2 mM L-Glutamine^13^. Unattached cells were removed 6-8 h after plating and the attached cells were cultured for 12-18 h to allow for full spreading out before performing efferocytosis assay^13^.

#### Murine splenic macrophages

Spleen single cell suspensions were obtained by grinding and filtering through 70 µM cell strainers^52^. F4/80^+^ macrophages were captured and purified using magnetic separation of F4/80^+^ cells by positive selection (Fisher Scientific, 50-112-5082) following the manufacture’s protocol.

#### Human monocyte-derived macrophages (HMDMs)

Human monocyte-derived macrophages (HMDMs) were differentiated from peripheral blood mononuclear cells (PBMCs)^13^. PBMCs were isolated from buffy coats of anonymous, de-identified healthy adult volunteer donors purchased from the New York Blood Center by Ficoll density gradient centrifugation^13^. PBMCs were cultured in RPMI-1640 medium supplemented with 20% HI-FBS and 50 ng/mL human macrophage colony-stimulating factor (M-CSF) for 7–8 days^13,53^.

### Efferocytosis assays

#### Preparation of apoptotic cells (ACs) and fluorescent labeling

ACs were generated by exposing isolated mouse thymocytes to ultraviolet light (UV, 254-nm wavelength) irradiation as described^13,54,55^. Briefly, single-cell suspension of mouse thymi of C57BL/6J wild-type mice were isolated and seeded in 10-cm non-TC-treated dish at 1 x 10^7^ cells per mL in 8 mL PBS, and then irradiated for a total of 15 min. Cell suspensions were then incubated at 37°C with 5% CO_2_ for 2.5 h. Annexin V staining was performed to confirm >90% apoptosis induction.

Fluorescent labeling of ACs was performed as described^13^. For single staining, ACs were labeled with 2 µM Hoechst 33342 (Thermo Fisher Scientific, 62249) for 30 min or 10 µg/mL TAMRA (Thermo Fisher Scientific, C1171) for 25 min. For co-labeling with Hoechst 33342 and pHrodo-Red (Thermo Fisher Scientific, P36600), ACs were first stained by 2 µM Hoechst 33342 for 30 min, followed by staining with 20 ng/mL pHrodo-Red for 20 min^56^. After washing the ACs with a 10-fold volume of PBS (Corning, 21-040-CV), ACs are ready to be used for efferocytosis assays.

#### *In vitro* efferocytosis assay

As we described^13,56^, macrophages (BMDMs or PMs) were differentiated in 6-well non-TC-treated plates at a density of ∼1 × 10^6^ per well. Fluorescently labeled ACs were co-incubated with macrophages at a 5:1 AC-to-macrophage ratio for 1 h in 2 mL DMEM basal media supplemented with 10% HI-FBS at 37°C, 5% CO_2_. Macrophages were then washed with 1X PBS gently for 5 times to remove unbound ACs. For flow cytometry-based quantification, macrophages were lifted using CellStripper (Corning, 25-056-CI), a non-enzymatic cell dissociation solution, for live-cell analysis.

#### *In vivo* thymus efferocytosis assay

Mice were injected intraperitoneally with 200 µL PBS containing 250 µg dexamethasone as previously described^13^. 18 h after injection, mice were euthanized, and thymi were dissected and weighed. One lobe was immersed in optimal cutting temperature (OCT) compound and snap-frozen for immunofluorescence staining to determine efferocytosis *in situ*^13^. Frozen sections were stained with terminal deoxynucleotidyl transferase dUTP nick end labeling (TUNEL, Thermo Fisher Scientific, C10618) to label ACs and CD68 antibody (Abcam, ab53444, 1:50) to label macrophages. The ratio of macrophage-associated ACs to free ACs was calculated to represent the capability of efferocytosis by thymus macrophages^13^. The other lobe was disaggregated into single cells to count the total number of thymocytes and quantify the percentage of Annexin V^+^ ACs using Annexin V Cellular Apoptotic kit (Invitrogen, A23204) by flow cytometry.

#### *In vivo* peritoneal macrophage efferocytosis assay

Mice were injected intraperitoneally with 1 × 10^7^ TAMRA-stained apoptotic mouse thymocytes in 300 μL PBS. 15 min after injection, mice were euthanized, and peritoneal exudates were collected. The pelleted cells were blocked with CD16/32 (Biolegend, 101302, 1:200), and then stained with F4/80-APC antibody (Biolegend, 123116, 1:100) for 15 min on ice to label macrophages. The percentage of TAMRA^+^ PMs was determined by flow cytometry^13^.

### AC injection in mice

#### Repeated injections to induce autoimmunity-like responses

Exposure to large numbers of ACs was used to evoke immune responses in mice as previously described^57^. Thymocytes from C57BL/6J wild-type mice were UV-irradiated to induce apoptosis and 2 x 10^7^ ACs per mouse were injected intravenously via tail vein injection. The injections were performed weekly for 4 weeks, paused for 2 weeks, followed by 2 additional injections at week 7 and week 8. One week after the last injection, mice were euthanized to collect serum and kidneys to assess immune responses.

#### One-time injection to assess AC accumulation in different organs

To determine AC accumulation in different organs upon injection, ACs were labeled with 10 µg/mL TAMRA (Thermo Fisher Scientific, C1171) for 25 min and 2 x 10^7^ ACs per mouse were injected intravenously via tail vein injection^58^. 24 h or 48 h post injection, mice were euthanized to harvest spleen, liver, and kidney. Control groups were injected with sterile PBS and were euthanized 48 h post-injection. Single cell suspensions were prepared to determine the percentage of TAMRA^+^ cells by flow cytometry. Cell suspensions from the control groups were used to assess autofluorescence and to guide gating strategy.

### Assessment of autoimmunity-related phenotypes in mice

#### Organ weight

Spleen, kidney, and subiliac lymph node (LN) were dissected and weighed.

#### Serum alanine aminotransferase (ALT) and creatinine

Peripheral blood was collected to obtain serum by centrifuging at 1400 X *g* for 10 min at 4°C. ALT and creatinine were determined using a Heska Element DC5X Veterinary Chemistry Analyzer with 120 µL serum delivered on ice and analyzed by the diagnostic lab at the Institute of Comparative Medicine (ICM), Columbia University Irving Medical Center.

#### Serum autoantibodies

Serum antinuclear antibodies (ANA) and anti-dsDNA antibodies were determined using mouse antinuclear antigens Ig’s (Total A+G+M) ELISA kit (Alpha Diagnostics International, 5210) and mouse anti-dsDNA Ig’s (Total A+G+M) ELISA kit (Alpha Diagnostics International, 5110).

#### Pathological analysis of the kidney

Kidneys were excised. One kidney was immersion-fixed in 4% formaldehyde (Thermo Fisher Scientific, 28908), embedded in paraffin, and coronal sections cut at 3 µm thickness^25^. Hematoxylin and eosin (H&E) staining was performed to grade the degree of mesangial hypercellularity and endocapillary hypercellularity/leukocyte infiltration on a semiquantitative scale of 0, 1+, 2+, 3+. For the degree of mesangial hypercellularity, 0 = none; 1+ = mild; 2+ = moderate; 3+ = severe hypercellularity. For the degree of endocapillary hypercellularity/leukocyte infiltration, 1+ = involving <25% glomeruli; 2+ = involving 25-50% glomeruli; 3+ = involving >50% glomeruli. Grading was performed blindly by a renal pathologist (V.D.D.).

The other kidney was embedded in optimal cutting temperature (OCT) compound and snap frozen. Cryostat sections were cut at 3 µm for periodic acid-Schiff (PAS) staining and for indirect immunofluorescent staining. For PAS staining, frozen sections were fixed with 4% formaldehyde for 20 min at room temperature. Cross sections were imaged with the Leica SCN 400 Slide Scanner. Glomeruli along the longitudinal line and the transverse line of the cross section were identified. Glomerular area was quantified and presented as the mean area of 12-15 glomeruli per mouse using the Aperio ImageScope software (Leica, V12.4.6.5003)^25^.

For immunofluorescence staining^59,60^, frozen sections were fixed with 4% formaldehyde for 20 min at room temperature. Permeabilization using 0.3% Triton X-100 in PBS was followed by blocking with 5% BSA. Slides were washed with PBS-0.05% Tween-20 (PBST) for three to five times between each step. For C1q staining, primary antibody incubation was carried out overnight at 4°C using rabbit monoclonal to C1q (Abcam, ab182451, 1:50), followed by secondary antibody incubation for 1 h at room temperature using Alexa-Fluor 555-goat anti rabbit IgG (H+L) (Invitrogen, A21428, 1:200). For IgG or wheat germ agglutinin (WGA) staining, Alexa-Fluor 555-conjugated anti-IgG (Invitrogen, A21422, 1:200) or Alexa-Fluor 488-conjugated WGA (Invitrogen, W11261, 1:5000) were used. DAPI (Invitrogen, P36931, 1:5000) was used for nuclear staining. Negative controls were included using rabbit IgG isotype control (Cell Signaling Technology, 3900S, 1:50). Slides were mounted with antifade mountant (Invitrogen, P36930) and imaged using the ImageXpress Micro4 high-content microscopy with a Nikon Plan Apo λ 20x/0.75 objective lens (Molecular Device) or Nikon Ti-S Automated Inverted Microscope (Nikon). The mean fluorescence intensity (MFI) within the glomeruli area was quantified by analyzing 20-25 glomeruli from two sections per mouse using Image J 1.53u.

### Western blotting

Cells from one-well of 6-well plate, i.e., about 1 x 10^6^ cells, were harvested using Cell Stripper (Corning, 25-056-CI). The cell pellet was then lysed on ice for 30 min in 70 µL 1 x RIPA buffer (MilliporeSigma, 20-188) supplemented with protease inhibitor cocktail and phosphatase inhibitor cocktail^61^, followed by centrifugation at 12,000 x *g* for 10 min at 4°C. The supernatants were transferred to new tubes and mixed with 5X SDS sample buffer (250 mM Tris-HCl, pH 6.8, 20% SDS, 30% (v/v) Glycerol, 10 mM DL-Dithiothreitol (DTT), 0.05% (w/v) Bromophenol blue). Samples were loaded onto a 3-8% Tris-Acetate gel (Thermo Fisher Scientific, EA0378BOX) and then transferred to 0.45 µm PVDF membrane. Blocking was performed in 5% milk in TBST at room temperature for 1 h. Primary antibody incubation was carried out at 4°C overnight, followed by incubation with secondary antibodies at room temperature for 1 h. Primary antibodies used for this study include anti-WDFY3 (a gift from Ai Yamamoto lab, 1:1000), anti-caspase-1 (mouse, eBioscience, 14-9832, 1:1000), and anti-caspase-1 (human, Thermo Fisher Scientific, MA547253, 1:500). Secondary antibodies include HRP-conjugated goat-anti-rabbit IgG (Cell Signaling Technology, 7074, 1:5000) and HRP-conjugated goat-anti-rat IgG (Cell Signaling Technology, 7077, 1:5000). Housekeeping protein antibodies include HRP-conjugated rabbit-anti-GAPDH (Cell Signaling Technology, 3683S, 1:5000), HRP-conjugated rabbit-anti-β-Actin (Cell Signaling Technology, 5125S, 1:2000), and HRP-conjugated rabbit-anti-Vinculin (Cell Signaling Technology, 18799S, 1:2000). Target protein and the respective housekeeping protein are derived from the same membrane.

### T cell activation by flow cytometry

T cell activation was characterized in blood T cells and splenic T cells in mice as previous described^25^. Blood samples were collected into a blood collection tubes with sodium heparin. Red blood cells (RBCs) were lysed by using ice-cold RBC lysis buffer (Invitrogen, 50-112-9751) for 10 min on ice. White blood cells were pelleted, washed, and resuspended in ice-cold FACS buffer containing 2% HI-FBS, 5 mM EDTA, 1 mM sodium pyruvate, and 20 mM HEPES in PBS. Splenic cells were obtained by mincing the tissue through a 70 μm strainer followed by RBC lysis. Cell suspensions were stained with an antibody cocktail against CD45-APC-Cy7 (BD Pharm, 557659, 1:100), Ly6-C/G-PerCP-Cy5.5 (BD Pharm, 552093, 1:100), CD115-PE (Invitrogen, 12-1152-83, 1:100), TCRβ-PB (Biolegend, 109226, 1:100), CD44-PE-Cy7 (Invitrogen, 25-0441-82, 1:100), CD62L-APC (Invitrogen,17-0620-82, 1:100) or CXCR3-APC (Invitrogen, 17-1831-82, 1:100), CD8-FITC (eBioscience, 11-0083-85, 1:100) or CD4-FITC (Biolegend, 100406, 1:100). CD4^+^ T cells were recognized as CD45^+^Ly6-C/G^-^CD115^-^TCRβ^+^CD4^+^ or CD45^+^Ly6-C/G^-^CD115^-^TCRβ^+^CD8^-^. The CD8^+^ T cells as CD45^+^Ly6-C/G^-^CD115^-^TCRβ^+^CD8^+^. The active form of CD4^+^ and CD8^+^ T cells were identified as CD44^+^CD62L^-^ or CXCR3^+^. Agilent NovoCyte Flow Cytometers were used to analyze the percentage of T cell populations.

### Antigen presentation assays

#### *In vitro* antigen presentation assay

MHC-II and MHC-I mediated antigen presentation by BMDMs *in vitro* was assessed using ovalbumin (OVA) peptides and OT-II and OT-I transgenic T cells, as modified from previously published protocols^25,26,62^. CD4^+^ T cells or CD8^+^ T cells were isolated from spleen homogenates of OT-II mice or OT-I mice, respectively, using CD4 (Miltenyi Biotec, 130-117-043) or CD8a (Miltenyi Biotec, 130-117-044) MicroBeads. Isolated T cells were labeled with 5 μM carboxyfluorescein succinimidyl ester (CFSE)-Violet Proliferation Dye 450 (VPD450, BD Horizon, 562158) for 25 min. For MHC-II-mediated antigen presentation, BMDMs were pulsed with OVA_323-339_ peptide (Anaspec, 50-843-886) for 24 h, followed by co-culture with CD4^+^ OT-II T cells for 72 h. For MHC-I-mediated antigen presentation, BMDMs were pulsed with OVA_257-264_ peptide (Anaspec, 50-844-730) for 24 h, followed by co-cultured with CD8^+^ OT-I T cells for 72 h. T cell proliferation was assessed by measuring CFSE-VPD450 dilution in CD4^+^ T cells (CD4-APC, Invitrogen, 17-0041-81, 1:100) and CD8^+^ T cells (CD8a-FITC, Invitrogen, 11-0081-82, 1:100) using a NovoCyte flow cytometer, as each round of division corresponds to a reduction in CFSE-VPD450 fluorescence intensity, thereby serving as a marker for T cell proliferation and activation.

#### *In vivo* antigen presentation assay

To assess MHC-II mediated antigen presentation *in vivo*^25^, mice were immunized with 20 µg OVA_323-339_ by subcutaneous injection. 24 h after injection, mice received 2 x 10^6^ CD4^+^CD90.1^+^ OT-II T-cells labeled with 10 μM CFSE-VPD450. After 72 h, splenocytes were isolated and stained with CD4-APC and CD90.1-PE (Invitrogen, 12-0900-83, 1:100) antibodies and analyzed using a NovoCyte flow cytometer. CD4^+^CD90.1^+^ T cells with diluted CFSE-VPD450 indicate proliferation of CD4^+^ T cells derived from CD90.1^+^ OT-II donor mice.

### Assessment of efferocyte-T cell interaction

The method to assess efferocyte-T cell interaction to induced T cell activation was modified from a previously published protocal^36^. After UV irradiation to induce apoptosis, ACs were collected and centrifuged at 500 x *g*. The pelleted ACs were resuspended in PBS supplemented with 2% HI-FBS and 2 mM EDTA at 2 x 10^7^ / mL, and incubated at 37°C, 5% CO_2_ overnight in sterile tubes. After overnight incubation, 0.5 mL cell suspensions were added to BMDMs in one well of 6-well plate and co-incubated for 24 h. After 24 h co-incubation, AC cell suspensions were removed by washing. CD4^+^ T cells were isolated from spleen homogenates of mice using CD4 MicroBeads (Miltenyi Biotec, 130-117-043), followed by CFSE-VPD450 labeling, and then added to BMDMs for 72 h co-incubation. The dilution of CFSE was evaluated using a NovoCyte Flow Cytometer^26,63^.

### LC3 lipidation measurement by flow cytometry

BMDMs were incubated with TAMRA-labeled ACs at a 5:1 AC-to-BMDM ratio at 37°C, 5% CO_2_ for 1 h efferocytosis. Unbounded ACs were washed away. BMDMs were collected and resuspended in 300 μL cold PBS with 20 µg/mL digitonin and incubated on ice for 10 min to permeabilize cells and allow non-membrane bound LC3 to be removed from cells. Permeabilized BMDMs were then centrifuged for 5 min at 750 x *g*, followed by incubation with Alexa Fluor 488-conjugated anti-LC3A/B antibody (Cell Signaling Technology, 13082S, 1:50) diluted in cold washing buffer (1X DPBS, 2% HI-FBS, 5 mM EDTA, 20 mM HEPES and 1 mM sodium pyruvate) for 15 min on ice to stain the membrane-bound lipidated LC3-II within the cells^64^. After staining, macrophages were washed with cold washing buffer. Cell pellets were resuspended in washing buffer and analyzed by flow cytometry by gating the TAMRA^+^ BMDMs to quantify the MFI of LC3A/B.

### Pro-IL-1β measurement by flow cytometry

For the detection of pro-IL-1β in splenic macrophages, cells were fixed and permeabilized using BD Cytofix/Cytoperm Fixation/Permeabilization kit (BD Biosciences, 554714), stained with pro-IL-1β-PE (eBioscience, 12-7114, 1:50) and its isotype control (eBioscience, 12-4301, 1:50), and analyzed by flow cytometry^25^.

### Cytokine and chemokine measurement in serum or culture media

The 26-plex ProcartaPlex Mouse Cytokine & Chemokine Convenience Panel 1 was used to analyze 26 protein targets simultaneously in a single well of serum sample using Luminex xMAP technology (Thermo Fisher Scientific, EPXR260-26088-901)^65^. Serum IL-18 was determined using the mouse IL-18 Duoset ELISA kit (R&D Systems, DY762505)^25^.

To determine how efferocytosis affect IL-1β secretion^66^, ACs were co-incubated with BMDMs for 8 h with a 5:1 AC-to-BMDM ratio, i.e., 5 x 10^6^ ACs were added to one-well of 6-well plate of differentiated BMDMs, that generally contains ∼1 x 10^6^ BMDMs. Media were collected to determine secreted IL-1β using mouse IL-1β/IL-1F2 Quantikine ELISA kit (R&D Systems, MLB00C).

### siRNA-mediated gene silencing

Non-targeting siRNA control (si-Ctrl) and WDFY3-targeting siRNA (si-WDFY3) (Dharmacon) were obtained from Dharmacon^13^. Human PBMCs were seeded at 5 x 10^6^ per well of 6-well non-tissue-culture plate for differentiation to HMDMs in RPMI-1640 medium supplemented with 20% HI-FBS and 50 ng/mL human M-CSF. On day 5, HMDMs were transfected with 25 pmol siRNAs using 7.5 μL Lipofectamine RNAiMAX (Invitrogen) in 300 μL Opti-MEM. On day 7, the culture media were refreshed before all treatments. To induce inflammasome activation^67^, HMDMs were primed by 20 ng/mL lipopolysaccharide (LPS) (Sigma-Aldrich, L3129) for 3 hours followed by one-hour treatment of 2 mM ATP (Thermo Fisher Scientific, R0441). To determine how efferocytosis affect caspase-1 cleavage, ACs were co-incubated with HMDMs for 4 h with a 5:1 AC-to-HMDM ratio, i.e., 2.5 x 10^6^ ACs were added to one-well of 6-well plate of differentiated HMDMs, that generally contains ∼0.5 x 10^6^ HMDMs. Unbound ACs were washed away and HMDMs were collected to determine caspase-1 cleavage by Western blotting.

### Pakistan Genome Resource

The Institutional Review Board at the Center of Non-Communicable Disease (NIH registered IRB 00007048) approved the study. All participants provided informed consent. Whole Exome Sequencing and Whole Genome Sequencing from the Pakistan Genome Resource (PGR) have been previous described^17,18^. Associations with circulating levels of IL-18, IgG, ALT, AST (aspartate aminotransferase), and creatinine were analyzed using linear regression adjusting for age, sex, age^2^, age*sex, top 10 genetic principal components and whole genome regression correction using REGENIE^68^.

### RNA-sequencing sample preparation and data analysis

BM cells were isolated from 8-10-week-old Ctrl and hWDFY3^KI/KI^ mice (n = 5 mice per genotype) and differentiated into BMDMs. Total RNAs were extracted from day 7 BMDMs using the Quick-RNA Miniprep Plus kit (Zymo Research, R1057) and the assessment of RNA integrity was performed using an Agilent 2100 Bioanalyzer. With a minimum of 300 ng input RNA, strand-specific, poly(A) + libraries were prepared and sequenced at 20 million 100-bp paired-end reads per sample. Data analysis was performed as we previously described^13^. Genes with an absolute fold-change > 1.5 and false discovery rate-adjusted P value <0.05 were considered as differentially expressed.

### Statistical analysis

Statistical analyses were conducted using GraphPad Prism 8. The nonparametric Mann-Whitney test was used to compare two groups. For scenarios involving two independent variables (factors) with two or more groups, a two-way ANOVA was performed. Tukey’s post hoc test was applied to adjust for multiple comparisons. Data are presented as mean ± standard error of the mean (SEM). A P-value of 0.05 or less was considered statistically significant. The exact P values, as well as the number of independent experiments or biological replicates, are specified in figures and figure legends.

## Data and code availability

All data and materials used in the study are available to researcher for purposes of reproducing or extending these analyses upon reasonable request. Data for RNA-sequencing will be deposited in the Gene Expression Omnibus (GEO) repository. Code will be available via the Zhang lab GitHub repository.

## Acknowledgements

The authors’ research work has received funding from the National Institutes of Health (NIH) (R01HL151611, R01HL168174, P01HL172741) and the National Center for Advancing Translational Sciences (NCATS) (Irving Scholar Award through UL1TR001873), the Marjorie and Lewis Katz Scholar Award, and the M. Iréne Ferrer Scholar Award (to H.Z.), R35HL145228 (to I.T.), R01NS077111 and R01NS101663 (to A.Y.), the Russell Berrie Foundation Diabetes Scholar Program (to X.W.), and the American Heart Association Postdoctoral Fellowship 21POST829654 (to X.W.), the American Heart Association Postdoctoral Fellowship 20POST35130003 and Career Development Award 23CDA1052177 (to F.L.), and an National Science Foundation (NSF) predoctoral fellowship (to K.R.C).

We would like to acknowledge the NIH funding sources to the Columbia Center for Translational Immunology (CCTi) Flow Cytometry Core by grant number S10OD020056 and S10RR027050 and P30DK063608; the NIH/NCI Cancer Center Support Grant P30CA013696 for the use of resources at the Columbia Genome Center; the Columbia Stem Cell Initiative (CSCI) Flow Cytometry Core under the leadership of Michael Kissner; and Nivia M. Urena at the Institute of Comparative Medicine (ICM) for technical support on blood chemistry analyzer. Schematic figures were created with BioRender.com.

## Author contributions

X.W. and H.Z. conceived and designed the project. X.W. performed majority of the experiments. Z.W. and J.C. assisted experiments. K.R.C., F.L., and R.K.S. provided technical support. X.W. and H.Z. performed bioinformatic analysis of the RNA-seq data. V.D.D. performed pathological assessment of kidney sections. K.S. and D.S. contributed to analyses using the PGR data. A.Y. established the mouse models for Cre-mediated and H.Z. drafted the manuscript. I.T. and A.Y. critically read the manuscript. H.Z. supervised the project and funding. All the authors have read the manuscript and provided input to the manuscript.

