## Supplemental Figures for "Macrophage WDFY3, a protector against autoimmunity"

### Extended figures

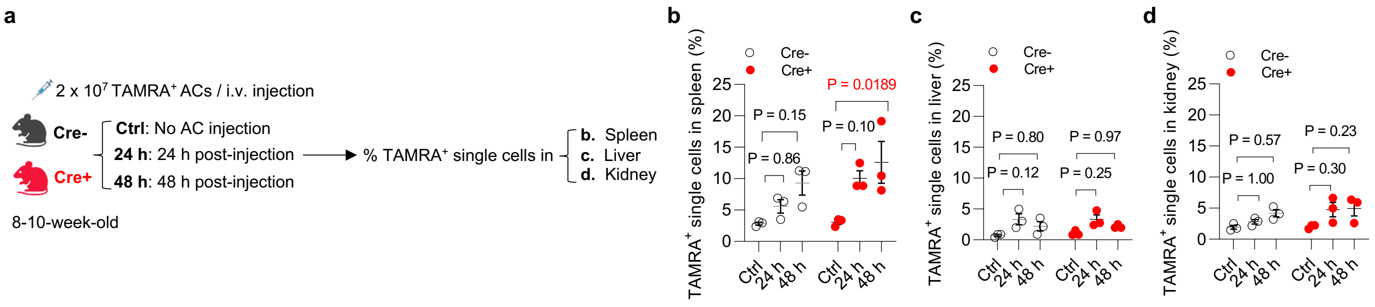


**Extended Data Fig. S1 | Accumulation of apoptotic cells (ACs) after intravenous injection. a**. Schematics of study design: 2 × 10$^7$ TAMRA-labeled apoptotic murine thymocytes (ACs) were injected intravenously via tail vein and mice were euthanized either 24 h or 48 h post-injection. Controls were injected with PBS and mice were euthanized 48 h post-injection. Spleen, liver, and kidney were dissected to determine the % of TAMRA$^+$ single cells that indicates AC accumulation by flow cytometry. TAMRA$^+$ cells were mostly accumulated in the spleen (**b**), but not liver (**c**) or kidney (**d**). Signals in controls with PBS injection (no ACs) represent autofluorescence. n = 3 Cre$^-$ vs 3 Cre$^+$ mice. Data are shown as mean ± SEM.

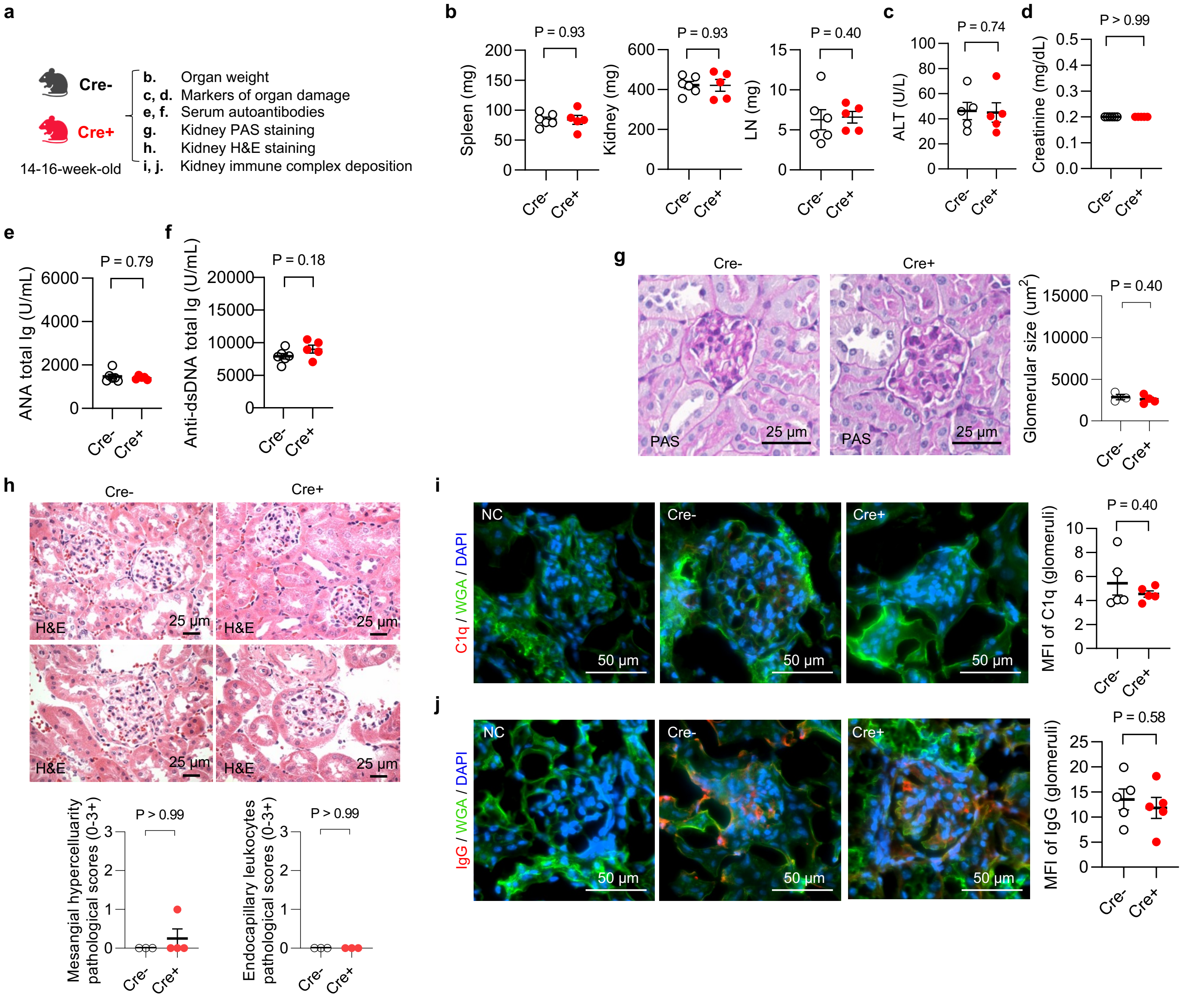


**Extended Data Fig. S2 | Myeloid knockout of *Wdfy3* does not affect autoimmune phenotypes in young mice. a,** Schematics of study design. Cre⁻ and Cre⁺ mice were euthanized at 14-16-week-old. Serum samples were collected to determine markers of organ damage and autoantibodies. Kidneys were dissected and sectioned for periodic acid-Schiff (PAS), H&E, and immunofluorescence staining. **b,** Organ weight was measured. n = 6 Cre⁻ vs 5 Cre⁺ mice. **c,** Serum alanine transaminase (ALT), a marker of liver damage. n = 5 Cre⁻ vs 5 Cre⁺ mice. **d**, Serum creatinine, a marker of kidney damage. n = 6 Cre⁻ vs 5 Cre⁺ mice. **e**, Serum anti-nuclear antibodies (ANA). n = 6 Cre⁻ vs 5 Cre⁺ mice. **f**, Serum anti-dsDNA antibodies. n = 6 Cre⁻ vs 5 Cre⁺ mice. **g**, PAS staining of kidney sections highlight glomerular basement membrane for the quantification of glomerular size. n = 3 Cre⁻ vs 4 Cre⁺ mice. **h**, H&E stain showing normocellular glomeruli in both Cre⁻ and Cre⁺ mouse kidneys. n = 3 Cre⁻ vs 4 Cre⁺ mice. **i** and **j,** Immunofluorescence staining of kidney sections using anti-C1q and anti-IgG antibodies. n = 5 Cre⁻ vs 5 Cre⁺ mice. Mean fluorescent intensity (MFI) in glomeruli were quantified. WGA, wheat germ agglutinin. Data are shown as mean ± SEM.

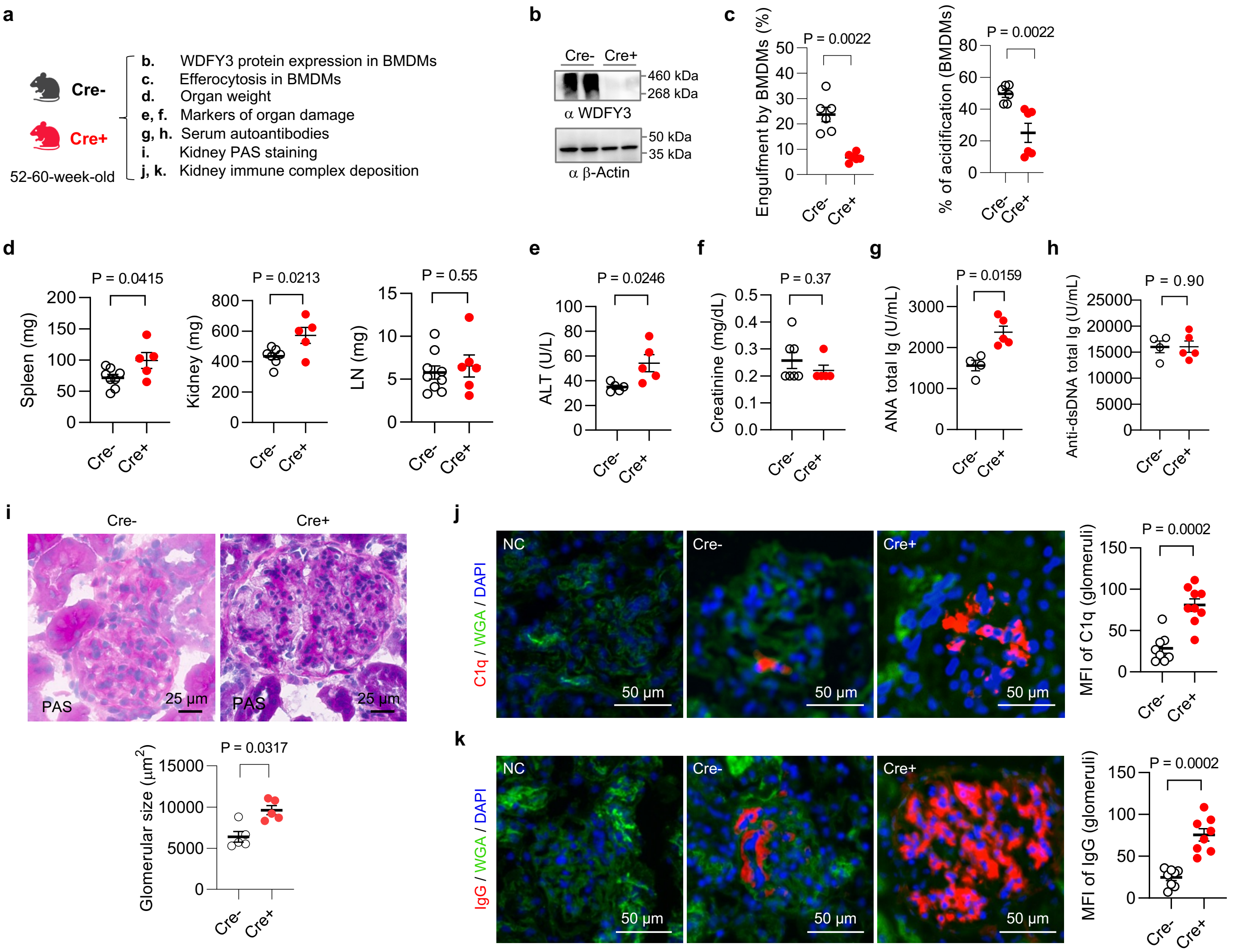


**Extended Data Fig. S3 | Myeloid knockout of *Wdfy3* exacerbates spontaneous autoimmunity in middle-aged mice. a,** Schematics of study design. Cre$^-$ and Cre$^+$ mice were euthanized at 52-60-week-old to assess efferocytosis and autoimmune phenotypes. Organ weight was measured. Serum samples were collected to determine markers of organ damage and autoantibodies. Kidneys were dissected for PAS and immunofluorescence staining. **b,** Successful knockout of *Wdfy3* at protein level was confirmed by Western blotting. **c,** BMDMs of Cre$^-$ mice showed impaired engulfment and lysosomal acidification of the engulfed ACs as determined by flow cytometry. n = 6 Cre$^-$ vs 6 Cre$^+$ mice. **d,** Organ weight. n = 8 Cre$^-$ vs 5 Cre$^+$ mice in spleen (left), n = 7 Cre$^-$ vs 6 Cre$^+$ mice in kidney (middle), and n = 9 Cre$^-$ vs 6 Cre$^+$ mice in lymph node (LN) (right). **e,** Serum ALT. n = 5 Cre$^-$ vs 5 Cre$^+$ mice. **f,** Serum creatinine. n = 7 Cre$^-$ vs 5 Cre$^+$ mice. **g,** Serum ANA. n = 4 Cre$^-$ vs 5 Cre$^+$ mice. **h,** Serum anti-dsDNA antibodies. n = 4 Cre$^-$ vs 5 Cre$^+$ mice. **i,** PAS staining of kidney sections for the quantification of glomerular size. n = 5 Cre$^-$ vs 5 Cre$^+$ mice. **j** and **k,** Immunofluorescence staining of kidney sections using anti-C1q and anti-IgG antibodies was performed. Mean fluorescent intensity (MFI) of C1q and IgG in glomeruli were quantified. WGA, wheat germ agglutinin. n = 8 Cre$^-$ vs 9 Cre$^+$ mice (j); n = 8 Cre$^-$ vs 8 Cre$^+$ mice (k). Data are shown as mean ± SEM.

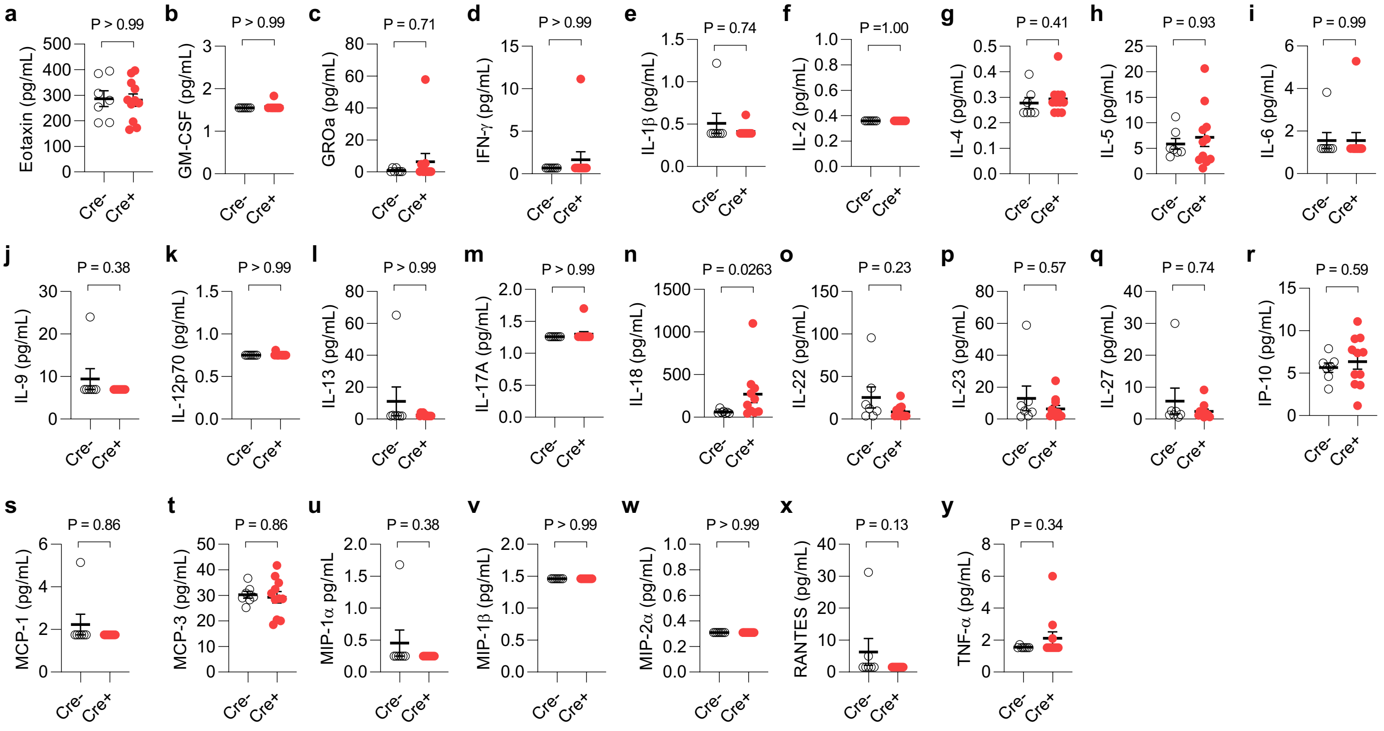


**Extended Data Fig. S4 | Serum cytokine and chemokine profiling.** Serum from Cre$^-$ or Cre$^+$ mice at 8-week-old were collected and assessed using a 26-plex ProCartaPlex mouse cytokine and chemokine panel. n = 7 Cre$^-$ vs 11 Cre$^+$ mice. Data are shown as mean ± SEM.

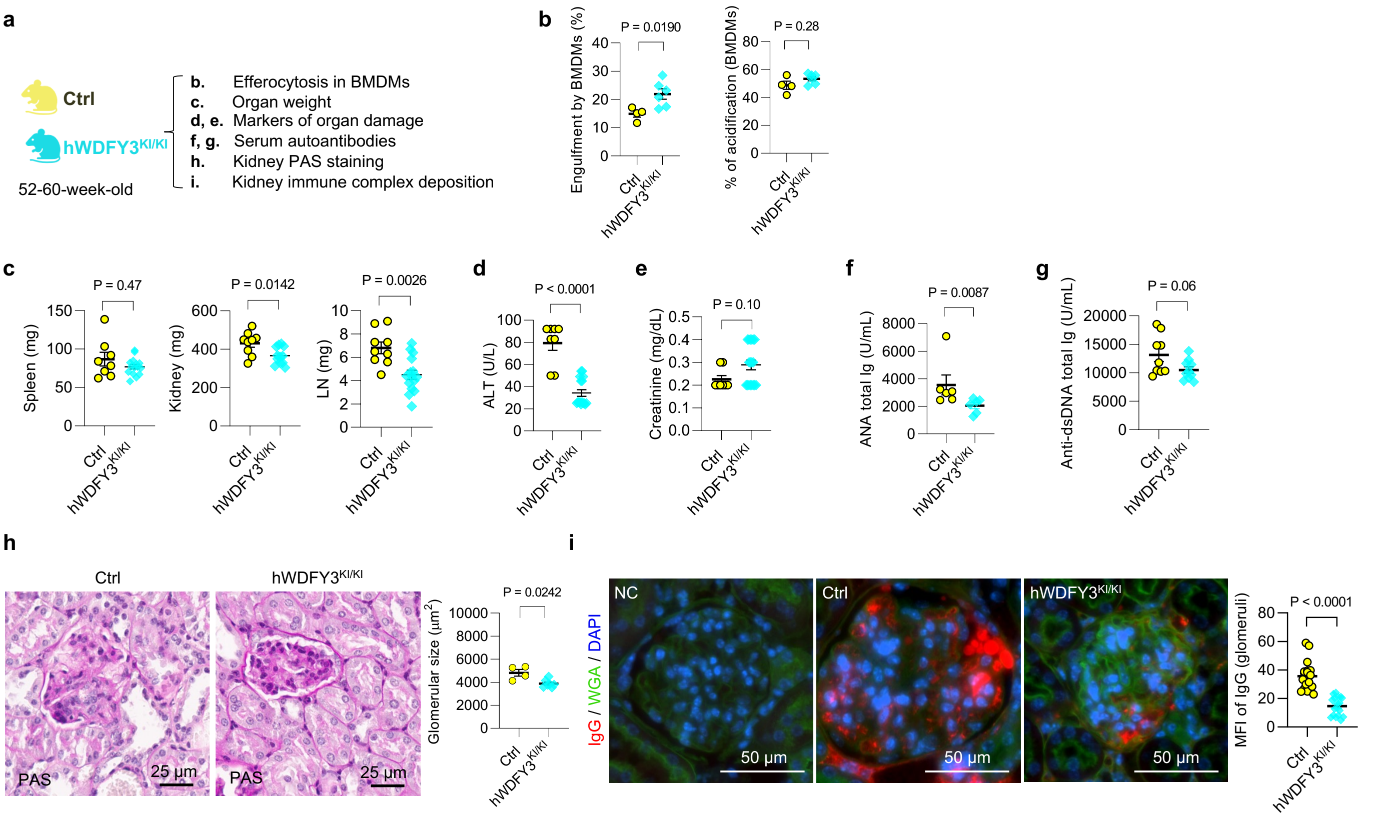


**Extended Data Fig. S5 | Overexpression of *WDFY3* mitigates spontaneous autoimmunity in middle-aged mice. a,** Schematics of study design. Age- and sex-matched Ctrl and hWDFY3$^{KI/KI}$ mice were euthanized at 52-60-week-old to assess efferocytosis, organ weight, markers of organ damage and autoantibodies in the serum, and kidney histology and immunocomplex deposition by PAS and immunofluorescence staining. **b,** Efferocytosis was enhanced in BMDMs of hWDFY3$^{KI/KI}$ mice. n = 4 Ctrl vs 9 hWDFY3$^{KI/KI}$ mice. **c,** Organ weight. n = 8 Ctrl vs 15 hWDFY3$^{KI/KI}$ mice in spleen (left), n = 9 Ctrl vs 12 hWDFY3$^{KI/KI}$ mice in kidney (middle), and n = 9 Ctrl vs 15 hWDFY3$^{KI/KI}$ mice in LN (right). **d,** Serum ALT. n = 8 Ctrl vs 16 hWDFY3$^{KI/KI}$ mice. **e**, Serum creatinine. n = 8 Ctrl vs 18 hWDFY3$^{KI/KI}$ mice. **f**, Serum ANA. n = 6 Ctrl vs 6 hWDFY3$^{KI/KI}$ mice. **g**, Serum anti-dsDNA antibodies. n = 9 Ctrl vs 10 hWDFY3$^{KI/KI}$ mice. **h**, PAS staining of kidney sections for the quantification of glomerular size. n = 4 Ctrl vs 7 hWDFY3$^{KI/KI}$ mice. **i,** Immunofluorescence staining of kidney sections using anti-IgG antibodies. n = 18 Ctrl vs 18 hWDFY3$^{KI/KI}$ mice. Data are shown as mean ± SEM.
